# CBL1/9-CIPK6 complex negatively regulates Respiratory burst oxidase homolog D in *Arabidopsis thaliana*

**DOI:** 10.1101/2025.03.30.646247

**Authors:** Niraj Kumar Vishwakarma, Nidhi Singh, Atish Sardar, Megha Choudhary, Debasis Chattopadhyay

**Affiliations:** National Institute of Plant Genome Research, Aruna Asaf Ali Marg, New Delhi, India

## Abstract

Plant innate immune response must be a well-balanced process with positive and negative regulations for the plants to survive. Calcium signalling is essential for pathogen-associated molecular pattern (PAMP)-driven Respiratory burst oxidase homolog D (RbohD)-mediated reactive oxygen species (ROS) burst. We show Calcium-sensors Calcineurin B like protein 1 (CBL1) and CBL9 and their interacting protein kinase CIPK6 negatively regulate RbohD activity and immune response in *Arabidopsis thaliana*. *Arabidopsis* mutant *cbl1cbl9*, like *cipk6*, exhibited enhanced resistance and ROS production when infected with the bacterial pathogen *Pseudomonas syringae* pv. *tomato* (*Pst*). CBL1 and CBL9 enhanced kinase activity of CIPK6. Plasma membrane localization of CBL1 and CBL9 and CIPK6 kinase activity were associated with the ROS production and immune response. CBL1/9-CIPK6 module interacts with RbohD at the plasma membrane and phosphorylates it’s N-terminal cytoplasmic domain at a non-conserved (Ser^33^) and a conserve (Ser^39^) Serine residue. While Ser^39^ phosphorylation increased RbohD activity, Ser^33^ phosphorylation drastically reduced it and superseded the effect of Ser^39^ phosphorylation. Replacement of Ser^33^ with alanine or aspartic acid made RbohD a super-active or low-active enzyme, respectively. Our study reports a direct mechanism of negative regulation of ROS production and plant immune response by a Calcium-signalling module in *Arabidopsis*. Overall, this study provides a novel insight into how calcium signaling integrates with immune regulation to prevent excessive ROS accumulation, ensuring a balanced plant immune response.

**One Sentence Summary:** Inactivation of RbohD by CBL1/CBL9-CIPK6 complex negatively regulates the plant immunity against *Pst* DC3000 infection in *Arabidopsis*.

## Introduction

Plants show resistance to microorganisms for their innate ability to recognize potential invading pathogens and to mount successful defences. Successful pathogens cause disease as they evade recognition or suppress host defence mechanisms or both (Staskawicz BJ, 2001). Various strains of the gram-negative bacterial pathogen *Pseudomonas syringae* have been used as models for understanding plant-pathogen interactions (Katagiri et al., 2002). Since the finding that *P. syringae* pathovar *tomato* (*Pst*) strain DC3000 (*Pst* DC3000) has the ability to infect *Arabidopsis* (*Arabidopsis thaliana*) in the laboratory in addition to its natural host tomato, it is being widely used to study not only the mechanism of plant immunity but also as a surrogate for studying mechanism of various effector proteins from other pathogens (Dong et al., 1991). *P. syringae* enters the host through wounds and stomata and multiply in the susceptible host in the intercellular spaces and depending on strains, ultimately shows necrosis surrounded by diffused chlorosis. On the other hand, in the resistant hosts, it triggers localized hypersensitive response (HR), a defence associated cell death (Dangl et al., 1996; Hammond-Kosack and Jones, 1997). HR is associated with the release of reactive oxygen species (ROS) and provides resistance against biotrophic pathogens. It is generally less effective against and sometimes beneficial to the necrotrophs, which complete their cell cycle in the dead tissues. In the cases of the hemibiotrophic pathogens such as, *Pst* DC3000 for *A. thaliana*, HR may be beneficial to the host at the beginning but may be harmful late in the interaction (Jupe et al., 2013; Münch et al., 2008).

The first line of plant defence against pathogen is the recognition of pathogen or the pathogen associated molecular patterns (PAMP) using the transmembrane pattern recognition receptors (PRRs). Pattern recognition leads to induction of PAMP-triggered immunity (PTI) and associated release of ROS (Asai et al., 2002; Boudsocq et al., 2010). Many bacteria inject various effector or avirulence (Avr) proteins in the host cells to interfere with PTI (Alfano and Collmer, 2004; Torres et al., 2006; He et al., 2006). Recognition of the effector proteins by the host-derived resistance (R) proteins leads to induction of effector triggered immunity (ETI) and release of the second burst of ROS for a prolonged period and sometimes HR-mediated cell death (Belkhadir et al., 2004; Nimchuk et al., 2003; Jones and Dangl, 2006; Alfano and Collmer, 2004; Torres et al., 2006; He et al., 2006). HR is a tightly regulated process and should be activated only when required and, therefore, is controlled at multiple layers (Goodman et al., 1994; Chandra-Shekara et al., 2006; Zhou and Zeng, 2017). Production of ROS in the apoplast is mediated by NADPH oxidases encoded by the respiratory burst oxidase homolog (Rboh) gene family. *Arabidopsis* genome encodes ten Rboh family genes. Many *Rboh* genes are transcriptionally upregulated by pathogen infection. Mutants lacking *RbohD* and *RbohF* genes exhibited very low ROS production when challenged with pathogens, indicating Rboh activity is required for pathogen-induced ROS production in the apoplast (Torres and Dangl, 2005).

Rapid increase in cytosolic calcium level ([Ca^2+^] _cyt_) is necessary for immune response (Grant et al., 2000; Lecourieux et al., 2006). Direct involvement of Ca^2+^ in ROS production by Rboh has been established. PTI-mediated rapid Ca^2+^ influx in the cytosol leads to phosphorylation and conformational changes in the N-terminal EF hands of RbohD (Ogasawara et al., 2008; Dubiella et al., 2013). Calcium dependent protein kinases StCDPK4 and StCDPK5 of potato, a receptor-like cytoplasmic kinase BIK1 and a cysteine-rich receptor like kinase CRK2 phosphorylate RbohB/D to enhance ROS production (Kobayashi et al., 2007; Kadota et al., 2014; Fujimoto et al., 2020; Li et al., 2014). Negative regulation of Rboh was also demonstrated with PBS1-like kinase 13 (PBL13), a negative regulator of PTI phosphorylates RbohD at the C-terminus resulting in increased ubiquitin-mediated degradation of RbohD and reduction of ROS production (Lee et al., 2020).

*Pst* DC3000 secretes various effectors, including AvrPto and AvrPtoB, which are recognised by Pto, a serine/threonine protein kinase (Cheng et al., 2011; Shan et al., 2008). A tomato (*Solanum* lycopersicum) variety resistant to *Pst* DC3000 triggers localized HR-mediated programmed cell death (PCD) upon recognition of AvrPto and AvrPtoB by Pto kinase as an effector-triggered immune response. Components of Ca^2+^-signalling pathways in tomato, Calcineurin B like protein 10 (SlCBL10) and its interacting protein kinase CBL-interacting protein kinase 6 (SlCIPK6) positively regulated ROS production and PCD (de la Torre et al., 2013). In contrast, AvrPto, the Avr protein in *Pst* DC3000 does not trigger ETI, rather elicit only PTI in *Arabidopsis* (Hauck et al., 2003). We have previously shown that CIPK6 (*Arabidopsis* CIPK6) functions as a negative regulator of immunity in *Arabidopsis*. The *cipk6* mutant showed reduced bacterial growth and higher hydrogen peroxide (H_2_O_2_) release as compared to the wild type plants (Sardar et al., 2017). In this report, we showed CBL1 and CBL9, the two CIPK6-interacting proteins also function together as the negative regulators of immune response to *Pst* DC3000 infection. Further, we showed that CBL1/9-CIPK6 complex interacts with RbohD at the plasma membrane and phosphorylates its N-terminus to inhibit its activity and negatively regulate the host immune response demonstrating a direct role of a calcium-mediated pathway in negative regulation of ROS production during plant immune response.

## Results

### *Arabidopsis* CBL1 and CBL9 interact with CIPK6 and negatively regulate immunity

Calcium signaling plays a crucial role in plant immunity by modulating defense responses to pathogen attacks. The CBL-CIPK signaling network, which consists of Calcineurin B-Like (CBL) proteins and their interacting CBL-Interacting Protein Kinases (CIPKs), has been implicated in various stress responses. In tomato, SlCBL10 has been shown to interact with SlCIPK6, forming a two-component signaling complex that regulates Effector-Triggered Immunity (ETI) and Programmed Cell Death (PCD) (de la Torre et al., 2013; Ren et al., 2021). However, whether a similar interaction exists in Arabidopsis and which CBLs participate in CIPK6-mediated immune signaling remain unknown.

To address this knowledge gap we performed a yeast-two-hybrid screen using Arabidopsis CIPK6 and all ten CBLs to identify potential interacting partners. Our experiment showed that CBL1, CBL2, CBL3, CBL4 and CBL9 were able to interact with CIPK6 in the yeast-two-hybrid assay and among these, CBL1 and CBL9 showed stronger interactions (Figure 1A and Supplementary Figure 1). Unlike in tomato, *Arabidopsis* CBL10 did not interact with CIPK6 in this assay. Expression of YFP-fused proteins in *N. benthamiana* leaves showed both the CBL1 and CBL9 proteins localize at the plasma membrane which was supported by western blot of subcellular protein fractions (Figure 1B and Supplementary Figure 2). Bimolecular fluorescence complementation (BiFC) experiment demonstrated *in planta* interaction between CIPK6 and CBL1/CBL9 in the plasma membrane while, CBL10 did not show any interaction in BiFC with CIPK6 of *Arabidopsis* (Figure 1C). CIPK6-CBL1/CBL9 interaction was also tested by expressing the CIPK6-YFP protein and 10xcMyc-CBL1/CBL9 proteins together in *N. benthamiana* leaves and co-immunoprecipitation with anti-GFP antibody followed by western blot with anti-cMyc antibody (Figure 1D). Addition of bacterially expressed CBL1 enhanced kinase activity of CIPK6 (Figure 1E), we investigated the involvement of CBL1 or CBL9 in host immune response. The *Arabidopsis* T-DNA insertion mutants *cbl1-/-* (*cbl1*) and *cbl9-/-* (*cbl9*) which lack CBL1 or CBL9 expression (Supplementary Figure 3B), respectively, did not show any significant difference in bacterial growth at three days after infiltration (3 dpi) with *Pst* DC3000 (Supplementary Figure 3A). *Arabidopsis* CBL1 and CBL9 are paralogous genes and their protein sequences show 97% similarity (Meena et al., 2015). CBL1 and CBL9 also showed overlapping functions in low potassium sensitivity (Xu et al., 2006; Cheong et al., 2007). Therefore, we infiltrated double mutant *cbl1-/-cbl9-/-* (*cbl1cbl9*) with *Pst* DC3000. The double mutant showed ten-fold less bacterial growth at 3 dpi as was observed in case of *cipk6* (Sardar et al., 2017). Restoration of CBL1 or CBL9 expression in the double mutant complemented the phenotype of the wild type plant in terms of bacterial growth, which is also substantiated by PTI-mediated H_2_O_2_ production (Figure 1F-G). *Pst* DC3000 triggers only PTI in *Arabidopsis*. Therefore, we repeated the same experiment with a *Pst* DC3000 strain expressing a bacterial effector AvrRPS4, which triggers both PTI and ETI in *Arabidopsis* (Sardar et al., 2017). Infiltration with *Pst* DC3000 (AvrRPS4) produced similar result of ten-fold less bacterial growth in *cbl1cbl9* (Supplementary Figure 4). Measurement of PTI- and ETI-mediated H_2_O_2_ production and the total salicylic acid (SA) content after *Pst* DC3000 (AvrRPS4) infiltration showed an inverse relationship with the bacterial growth (Supplementary Figure 5). The results described above showed that CBL1 and CBL9 redundantly act as the negative regulators of immune response in *Arabidopsis*.

**Figure 1.**
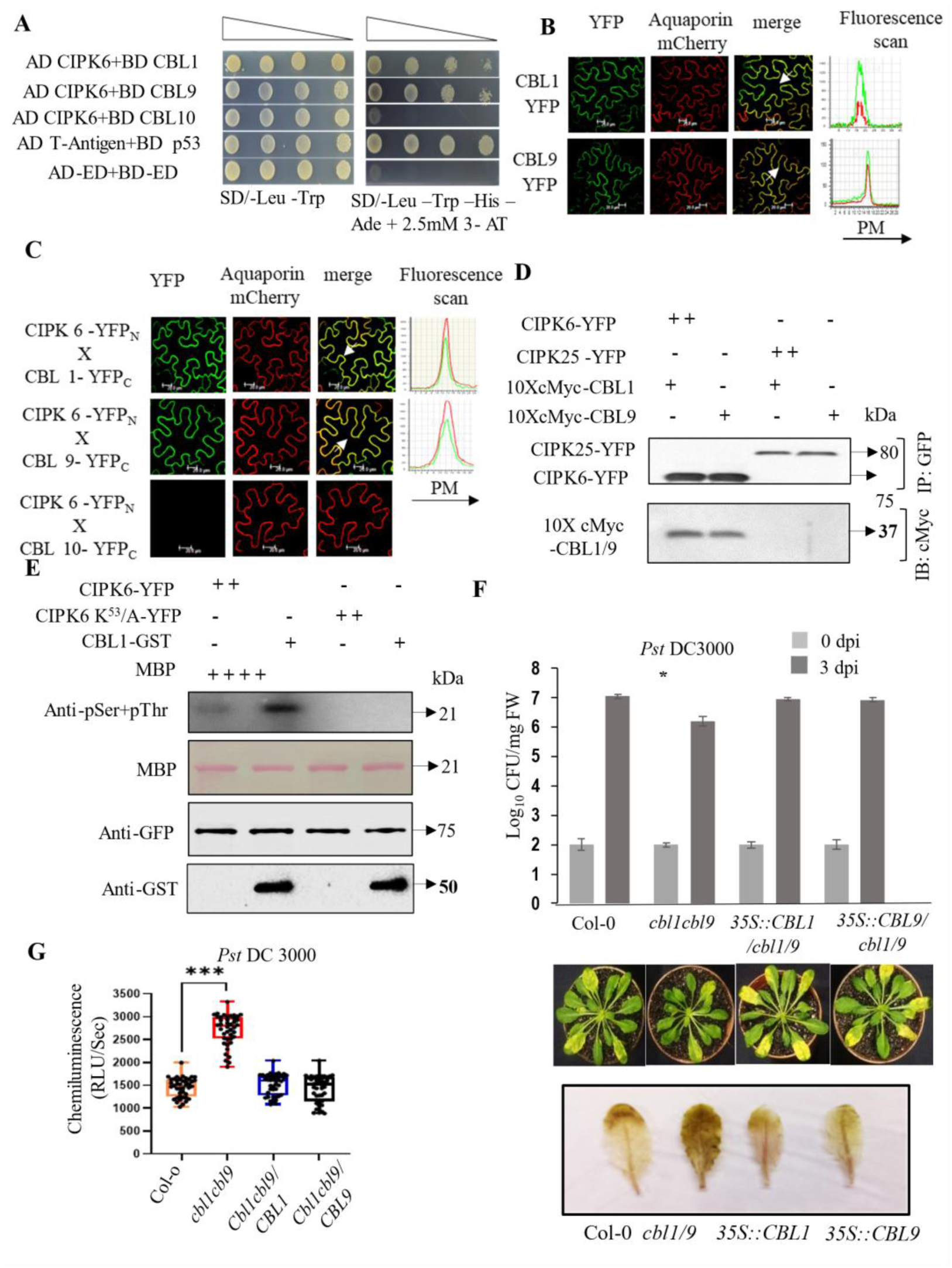
CBL1 and CBL9 localize, interact with CIPK6 in the plasma membrane (PM), promote its kinase activity, and negatively impact plant defense. (A) Yeast Two Hybrid (Y2H) assay show that CIPK6 interacts with CBL1 and 9 but not with the CBL10. Co-transformation of pGADT7 T-Antigen + pGBKT7-p53 and Empty bait vector + prey vector was used as positive and negative control, respectively. (B) Localization study shows that YFP tagged CBL1 and CBL9 (green) overlapped with PM marker Aquaporin-mCherry (red) and give yellowish appearance. (C) BiFC shows that CIPK6-YFP^N^ interacts with CBL1-YFP^C^ and CBL9-YFP^C^ at PM *in planta* in *Nicotiana benthamiana*. Fluorescence due to reconstitution of truncated YFP (green) merges with PM marker Aquaporin-mCherry (red) gives yellowish appearance. Infiltration of CBL10-YFP^C^ with CIPK6-YFP^N^ do not reconstitute YFP. CBL10-YFP^C^ does not interact with CIPK6-YFP^N^ and was used as negative control. (D) CIPK6-YFP and CIPK25-YFP with 10xcMyc-CBL1 or 10xcMyc-CBL9 were co-expressed in *N. benthamiana* leaves by agroinfiltration to perform the co-immunoprecipitation (Co-IP). *In vivo*, CIPK6 interacts with CBL1 and 9. No interaction was detected when YFP tagged CIPK25 was agro-infiltrated with 10xcMyc-CBL1 or 10xcMyc-CBL9. (E) *In vitro* kinase activity of CIPK6 is enhanced in presence of CBL1. Either in presence or absence of GST-CBL1, MBP (Myelin Basic Protein) substrate was incubated in kinase buffer with active CIPK6 and inactive kinase CIPK6 K^53^/A followed by immunoblotting with anti -phospho serine/threonine antibody. (F) *Pst* DC3000 (OD_600_ = 0.0005) was manually infiltrated into 4 week-old leaves of the Col-0, *cbl1cbl9*, *cbl1cbl9/CBL1* and *cbl1cbl9/CBL9*. Bacterial count was evaluated at 0 and 3 days post-infection (dpi). The asterisks indicate a significant difference following two-way ANOVA (α=0.05). (G) ROS burst measurement in Col-0, *cbl1cbl9*, *cbl1cbl9/CBL1* and *cbl1cbl9/CBL9*. Box-Plot showing the plot of maximum values of ROS attained during PTI. *n* > 35 for ROS burst. The asterisks indicate a significant difference following two-way ANOVA (α=0.05).

### Subcellular localization of CBL1 and CBL9 and their complex with CIPK6 affects their role in immune response

The subcellular localization of CBL-CIPK complexes plays a critical role in defining their functional outcomes in various plant signaling pathways. Previous studies have demonstrated that CIPK6 localization depends on its interacting CBL partners. In root cells, CIPK6 localizes to the cytosol and nucleus (Chen et al., 2013), where CBL1 and CBL4 show marginal expression (Klepikova et al., 2016). However, CBL1, CBL4, CBL5, and CBL9 possess a conserved MGCXXS motif, making them substrates for myristoylation and palmitoylation, which are essential for their proper targeting to the plasma membrane (Batistič et al., 2008). The N-myristoylation have been shown to be crucial for CBL1 and CBL9 function in salt stress responses, (Ishitani et al., 2000; Batistič et al., 2008), but their role in plant immunity remains unclear.

To explore the role of subcellular localization of CBL1/9-CIPK6 complex in immune response, we replaced the myristoyl-accepting second amino acid glycine of CBL1 and CBL9 with alanine. Subcellular localization of CBL1 G^2^/A and CBL9 G^2^/A was investigated by expressing the YFP-fused proteins in *N. benthamiana* leaves and by subcellular fraction followed by immunoblot. While the wild type CBL1 and CBL9 proteins localized at the plasma membrane, the CBL1 G^2^/A and CBL9 G^2^/A localized primarily in the cytoplasm and nucleus and marginally in the plasma membrane (Figure 2A-B and Supplementary Figure 6). CIPK6 interacted with CBL1 and CBL9 at the plasma membrane. However, CBL1 G^2^/A and CBL9 G^2^/A interacted with CIPK6 mostly at their altered location *i.e.,* in cytoplasm and nucleus and at a very low level at the plasma membrane (Figure 2C and Supplementary Figure 7). To explore the effect of altered subcellular location of CBL1- and CBL9 complexes in their role in immune response, we expressed CBL1 G^2^/A or CBL9 G^2^/A in *cbl1cbl9* line and compared the bacterial growth at 3 dpi with the mutant line expressing the wild type versions of these molecules.

**Figure 2.**
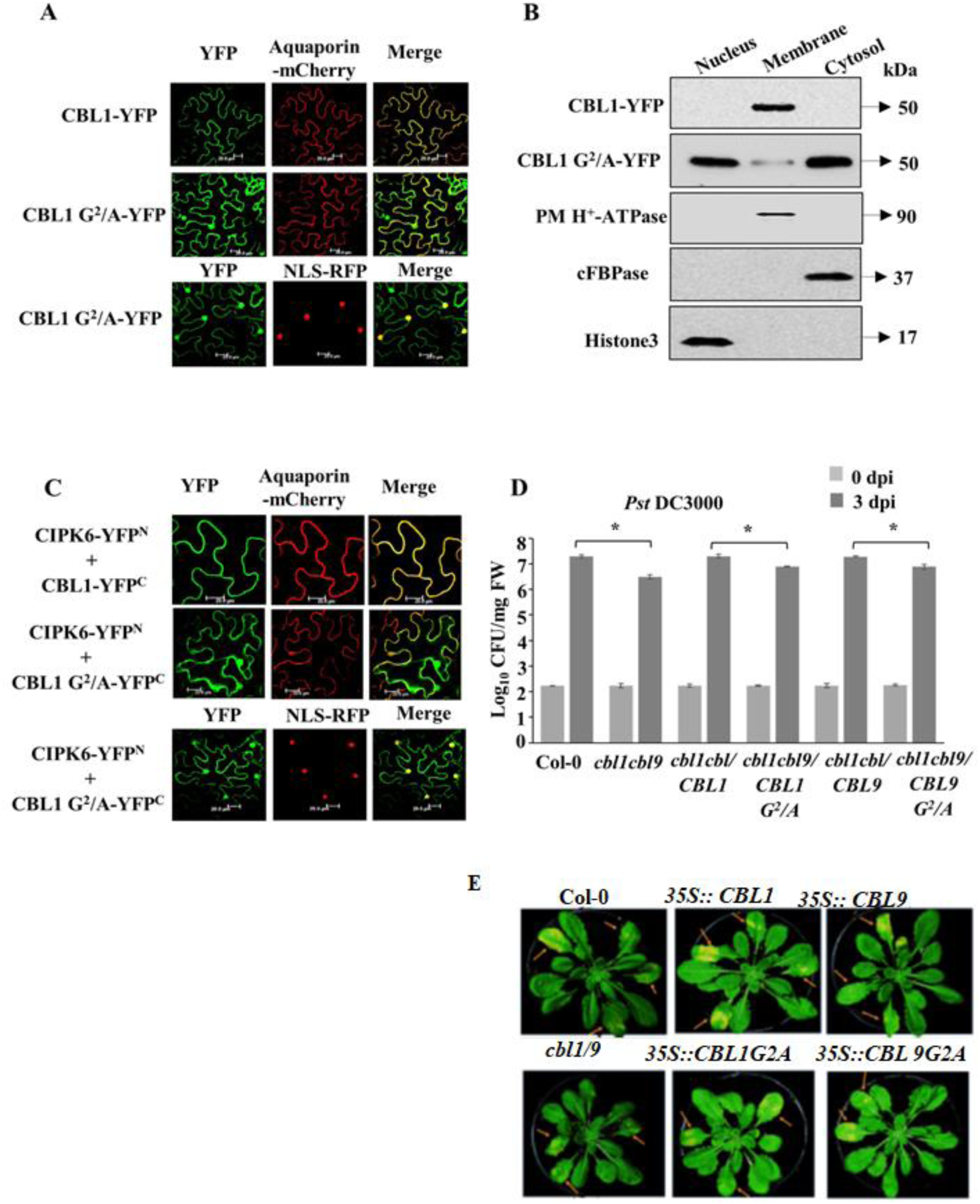
CBL 1 and CBL9 regulate the localization of CIPK6 and negatively affect plant immunity. **(A)** Mutation at myristylation site (G^2^/A) of CBL1 altered its localization from PM to cytosol and nucleus. YFP tagged CBL1 G^2^/A (Green) co-localized with nucleus marker NLS-RFP (red) and produced yellow color. YFP tagged CBL1 G^2^/A (Green) did not overlap with Plasma Membrane (PM) marker Aquaporin-mCherry (red). **(B)** Immunoblot of subcellular fractions of CBL1-YFP and CBL1 G^2^/A-YFP infiltrated *Nicotiana* leaves shows that CBL1 is exclusively localized at PM whereas CBL1 G^2^/A is predominantly present in nucleus and cytosol and just marginally present at PM. Immunoblot was probed with antibodies against GFP for CBL1 and CBL1 G^2^/A, plasma membrane H^+^ -ATPase, Cytosolic fructose-1,6-bisphosphatase (cFBPase), and H3 histone. **(C)** CBL1 G^2^/A-YFP^C^ interacted with CIPK6-YFP^N^ in nucleus and cytosol, *in planta*. Fluorescence due to reconstitution of truncated YFP (green) merged with nucleus marker NLS-RFP (red) produced yellow color. Fluorescence due to reconstitution of truncated YFP (green) did not overlap with PM marker Aquaporin-mCherry (red). **(D)** Bacterial load was determined in Col-0, *cbl1cbl9, cbl1cbl9/CBL1*, *cbl1cbl9/CBL1 G^2^/A*, *cbl1cbl9/CBL9* and *cbl1cbl9/CBL9 G^2^/A* at 0- and 3-days post-infection (dpi) of *Pst* DC3000 (OD_600_ = 0.0005). The asterisk sign indicates a significant difference between the marked genotypes, as observed through two-way ANOVA (α=0.05). * Localization of CBL9, CBLG^2^/A and their interaction with CIPK6 shown in Supplementary figure 6 and Supplementary figure 7.

While the expression of wild type CBL1 or CBL9 in *cbl1cbl9* line completely restored the bacterial growth in the wild type (Col-0) plants, *cbl1cbl9* lines expressing CBL1 G^2^/A and CBL9 G^2^/A variants lacking myristoylation sites were partially able to restore the bacterial growth as compared to the wild type plants or the *cbl1cbl9* plants expressing wild type versions of CBL1 or CBL9 by showing only ∼5-fold increase in the bacterial count as compared to the *cbl1cbl9* plant suggesting plasma membrane localization of CBL1 and CBL9 was required for their function in immune response (Figure 2D). Infiltration with avirulent strain *Pst* DC3000 (AvrRPS4) produced the similar result of partial restoration of activities of CBL1 and CBL9. The H_2_O_2_ peaks signifying PTI- and ETI-mediated immune response also followed the result obtained by the bacterial growth assay (Supplementary Figure 8B).

### *Pst* infection decreases CIPK6 expression and its kinase activity is associated with immune response

CBL-interacting Protein kinases (CIPKs) play a crucial role in plant stress responses, but their function in immunity remains complex and species-dependent. While previous studies have shown that *SlCIPK6* in tomato acts as a positive regulator of plant immunity by promoting H₂O₂ production and localized programmed cell death upon *Pst DC3000* infection (de la Torre et al., 2013), our findings showed that Arabidopsis *CIPK6* functions as a negative regulator of immunity in its native system. The *Arabidopsis cipk6-/-* (*cipk6*) mutant exhibited reduced bacterial growth and enhanced H_2_O_2_ generation with respect to the wild type plant in response to *Pst* DC3000 infection (Sardar et al., 2017). Consistent with its negative regulatory role in plant immunity, we observed a decline in CIPK6 gene expression up to five-folds at the twelfth hour after bacterial infiltration and then increased afterwards although not to the level before infiltration (Figure 3A). We cloned *Arabidopsis* CIPK6 cDNA fused to 10x-cMyc under its own promoter (P_CIPK6_) and expressed it in *cipk6* mutant to monitor CIPK6 protein expression upon *Pst* DC3000 infiltration. CIPK6 protein abundance quickly and gradually decreased to ∼2-fold within 90 min after infiltration. CIPK6 kinase activity also declined to ∼2-fold with 15 min after infiltration (Figure 3B, C and D).

**Figure 3.**
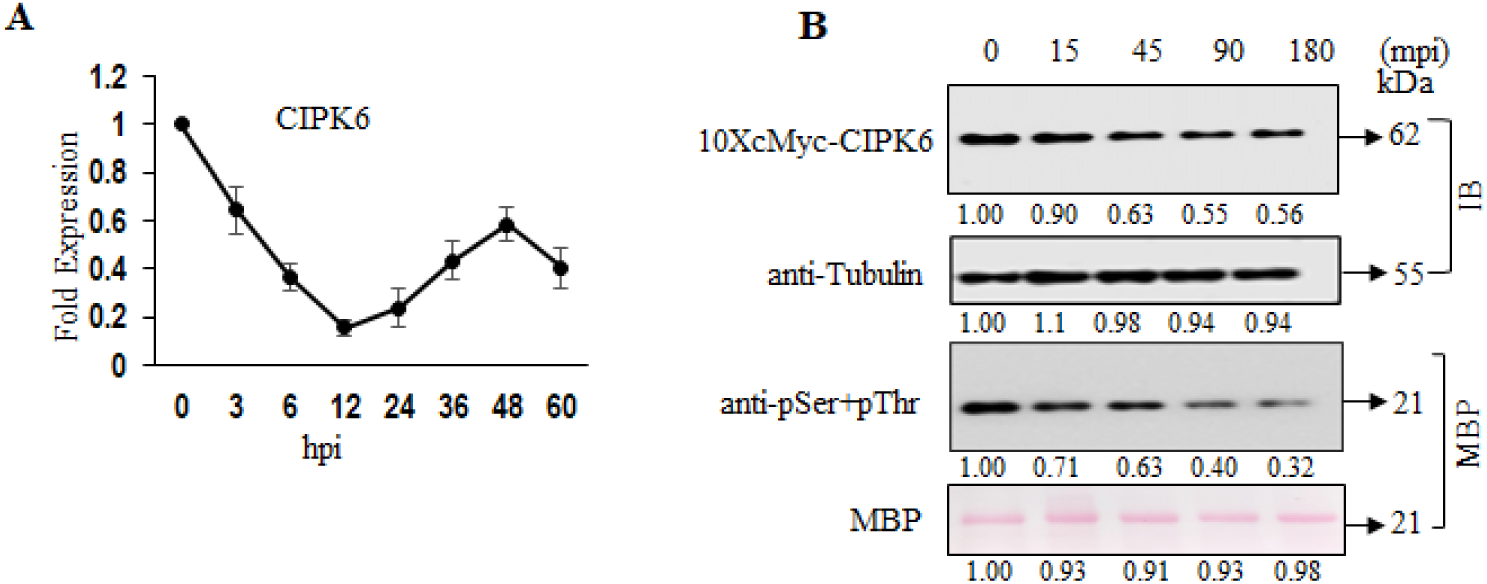
The transcript level and abundance of CIPK6 decreases over the time after pathogen inoculation. **(A)** Expression of *CIPK6* was assessed by qPCR in Col-0 after *Pst* DC3000 infection at the indicated time points. **(B)** Immunoblot shows that abundance of CIPK6 protein decreased gradually after *Pst* infection. *In vitro* kinase activity reflected that trans-kinase activity of CIPK6 also get declined due to gradual loss of CIPK6 protein after *Pst* infection. Numbers below represent the protein band intensity.

In order to investigate the role of the kinase activity of *Arabidopsis* CIPK6 in negatively regulating the immunity, we created truncated and mutated forms of CIPK6 with differential kinase activities. Following the previous reports, a conserved lysine (K^53^) in the kinase domain and a conserved threonine (T^182^) in the activation loop of CIPK6 were individually replaced with alanine and aspartic acid, respectively, to create CIPK6 K^53^/A and CIPK6 T^182^/D (Guo et al., 2001; Gong et al., 2002; Liu et al., 2000). The C-termini of CIPK proteins are known to autoinhibit their own kinase activities by blocking their kinase domains (Sánchez-Barrena et al., 2007). Therefore, a C-terminal deletion mutant was created by removing the C-terminus of CIPK6 T^182^/D (CIPK6 T^182^/DΔC). These constructs were expressed in *Escherichia coli* as Glutathione-S-transferase (GST)-fused proteins and the affinity-purified proteins were tested for *in vitro* autophosphorylation activity. Expectedly, CIPK6 K^53^/A did not show any autophosphorylation activity while, CIPK6 T^182^/D displayed a higher autokinase activity than the CIPK6. Autokinase activity of CIPK6 T^182^/DΔC was the highest among these three variants (Figure 4A and Supplementary Figure 9A). These constructs were expressed as yellow fluorescence protein (YFP)-fused proteins in *N. benthamiana* leaves and immunoprecipitated with GFP antibody to assess their substrate phosphorylation activities using myelin basic protein (MBP) as the substrate. Transphosphorylation activities of these three variants followed their autophosphorylation activities (Figure 4B and Supplementary Figure 9B). CIPK6 and its three variants having differential kinase activities were expressed in the *cipk6* mutant line and the growth of *Pst* DC3000 was assessed in the stable transgenic lines.

**Figure 4.**
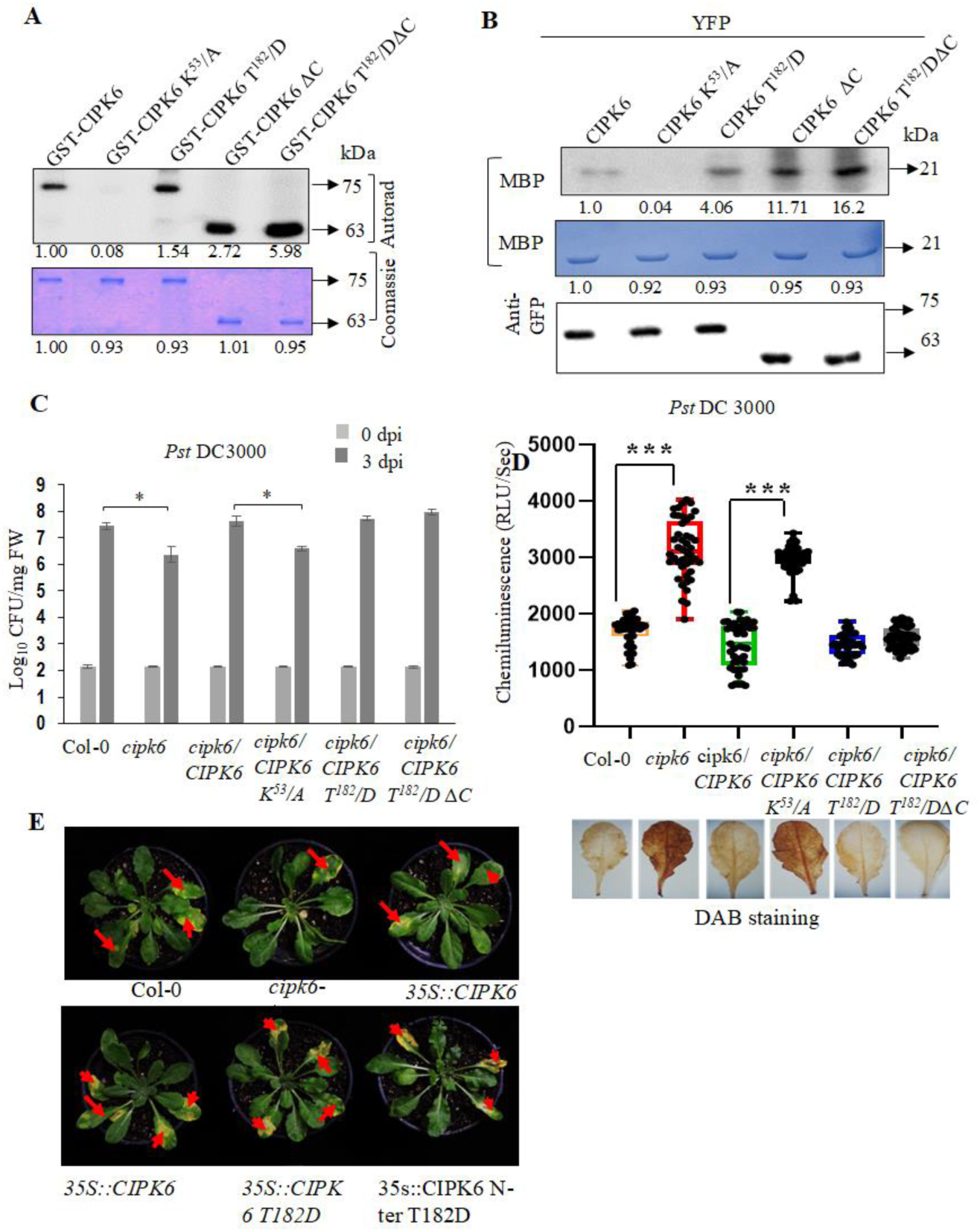
The CIPK6 kinase activity negatively controls plant immunity during pathogenesis. **(A)** To study autophosphorylation of CIPK6, recombinant GST-CIPK6 and its variants were incubated with kinase buffer (20 mM Tris-Cl, pH 8.0, 5 mM MnCl_2,_ 1 mM CaCl_2_, 0.1 mM EDTA, 1 mM DTT, and 10 µCi γ-^32^P ATP). In autophosphorylation assay, CIP6 K^53^/A (kinase dead) lacked kinase activity while CIPK6 exhibited limited auto-phosphorylation activity. CIPK6 variants CIPK6 T^182^/D, and CIPK6ΔC showed increased auto-phosphorylation activity and CIPK6 T^182^/DΔC (Super kinase) displayed the highest kinase activity (Upper Panel). SDS-PAGE gel was stained with Coomassie brilliant blue (lower panel). Numbers below show the protein band intensity. **(B)** CIPK6-YFP and its variant were transiently expressed in *N. benthamiana* and immunoprecipitated (IP) using anti-GFP antibody for transphosphorylation. For kinase assay, immunoprecipitated CIPK6 and its variant were incubated with MBP in kinase reaction buffer. Kinase reactions were resolved on 12% SDS-PAGE and autoradiographed (upper panel). Numbers below represent protein band intensity. Equimolar loading of CIPK6 and its mutant protein was confirmed by western blot using anti-GFP (middle panel). For equal loading of MBP, SDS-PAGE gel was stained with Coomassie brilliant blue (lower panel). **(C)** Bacterial growth was assessed in Col-0, *cipk6, cipk6/CIPK6*, *cipk6/CIPK6 K^53^/A*, *cipk6/CIPK6 T^182^/D*, and *cipk6/CIPK6 T^182^/DΔC* at 0 and 3 dpi of *Pst* DC3000 (OD_600_ = 0.0005). The asterisks above the indicated line show a significant difference observed by two-way ANOVA (α=0.05). **(D)** ROS burst measurement in Col-0, *cipk6, cipk6/CIPK6*, *cipk6/CIPK6 K^53^/A*, *cipk6/CIPK6 T^182^/D*, and *cipk6/CIPK6 T^182^/DΔC*. Box-Plot showing the plot of maximum values of ROS attained during PTI. *n* > 35 for ROS burst. The asterisks denote a significant difference after two-way ANOVA (α=0.05).

As reported before, *cipk6* showed ten-fold less bacterial growth as compared to the *Col-0* at 3 dpi, while restoration of CIPK6 expression in the mutant line resulted in abolition of resistance (Figure 4C and E). However, expression of CIPK6 T^182^/D, which showed a higher *in vitro* kinase activity did not show a further higher bacterial growth in comparison to the *cipk6*/CIPK6 line indicating a functional saturation of CIPK6. Expression of the inactive kinase variant CIPK6 K^53^/A in *cipk6* did not alter the bacterial growth in comparison to the mutant significantly, suggesting kinase activity of CIPK6 is required for the negative regulation of immune response. Expression of CIPK6 T^182^/DΔC, showing the highest *in vitro* kinase activity among the variants showed the highest bacterial growth and lowest immunity, but not significantly higher than the *cipk6*/CIPK6 line. PTI-mediated H_2_O_2_ release expectedly followed an inverse relation with the bacterial growth (Figure 4D and F). All the results described above demonstrated that kinase activity of *Arabidopsis* CIPK6 was required for its negative regulation of immune response.

### CBL1/9 and CIPK6 interact with RbohD and CIPK6 phosphorylates RbohD

Reactive oxygen species (ROS) play a critical role in plant immunity, acting as signaling molecules that modulate defense responses. Plasma membrane-localized NADPH oxidases, known as respiratory burst oxidase homologs (Rbohs), are key generators of ROS during pathogen attack. The activation of Rboh proteins is regulated by phosphorylation at their N-terminal cytoplasmic domains, a process mediated by Ca^2+^-dependent protein kinases (CPKs) and calcineurin B-like (CBL)-interacting protein kinases (CIPKs) (Kobayashi et al., 2007; Ogasawara et al., 2008; Dubiella et al., 2013). Previous studies have demonstrated that CBL1-CIPK26 and CBL1/9-CIPK26 complexes phosphorylate RbohC (Zhang et al., 2018) and CBL1/9-CIPK26 complexes phosphorylate RbohC and RbohF, respectively, enhancing ROS production in heterologous HEK293T cells (Drerup et al., 2013). However, the involvement of other CIPKs in Rboh phosphorylation remains largely unexplored. Given that tomato CIPK6-CBL10 has been shown to interact with RbohB (de la Torre et al., 2013), we sought to investigate whether Arabidopsis CIPK6 also interacts with and phosphorylates its ortholog, RbohD. Using YFP-tagged RbohD expressed in Nicotiana benthamiana, we confirmed its plasma membrane localization. BiFC assay by reconstitution of the fluorescent YFP protein showed interaction between CIPK6 and RbohD at the plasma membrane (Figure 5A). Kinase activity of CIPK6 was not essential for this interaction as CIPK6 K^53^/A also showed similar interaction (Supplementary Figure 10). As both the CBL1 and CBL9 localize at the plasma membrane and interact with CIPK6, we repeated the same experiment to explore interaction between RbohD and these CBLs. Both the CBL1 and CBL9 exhibited interaction with RbohD at the plasma membrane (Figure 5B). To substantiate the *in planta* interaction between RbohD, CIPK6 and CBL1, we expressed combination of RbohD-YFP, 10xcMyc-CIPK6, 3xFlag-CBL1 together in *N. benthamiana*. Immunoprecipitation of RbohD with anti-GFP antibody followed by immunoblot with anti-cMyc and anti-Flag antibody could detect presence of CIPK6 and CBL1 in an *in planta* complex with RbohD. However, no signal was detected when CIPK6 and CBL1 were replaced with CIPK25 and CBL10, respectively (Figure 5C). CIPKs are known as serine/threonine kinases (Hashimoto et al., 2012). To explore whether CIPK6 was able to phosphorylate RbohD, bacterially expressed GST-fused N-terminal cytoplasmic domain of RbohD protein (Kadota et al., 2014; Li et al., 2014) was used as a substrate to GST-CIPK6 and its kinase-inactive mutant CIPK6 K^53^/A. CIPK6, but not its kinase-inactive mutant was able to phosphorylate RbohD. The phosphorylation was detected with the anti-phosphoserine antibody but not with anti-phosphothreonine antibody, demonstrating *Arabidopsis* CIPK6 phosphorylates RbohD at serine residues (Figure 5D). Direct interaction between the bacterially expressed CIPK6-His and N-terminal domain of RbohD protein was detected by His-pull down followed by immunoblot (Figure 5E). Phosphorylated RbohD was analysed by mass spectrometry to detect the phosphorylation site. Four serine residues Ser^30^, Ser^33^, Ser^39^ and Ser^119^ of RbohD were detected to be phosphorylated by CIPK6 (Supplementary Figure 12). These residues were serially replaced with alanine and incubated with CIPK6 to confirm the phosphorylation sites. Replacement of Ser^30^ and Ser^119^ with alanine did not alter the phosphorylation intensity while individual replacement of Ser^33^ and Ser^39^ reduced phosphorylation intensity substantially while, Ser^33^ appeared to be the major CIPK6-mediated phosphorylation site. Replacement of both Ser^33^ and Ser^39^ with alanine completely abolished CIPK6-mediated phosphorylation of RbohD confirming Ser^33^ and Ser^39^ of RbohD are the target sites of CIPK6 (Figure 5F and Supplementary Figure 11).

**Figure 5.**
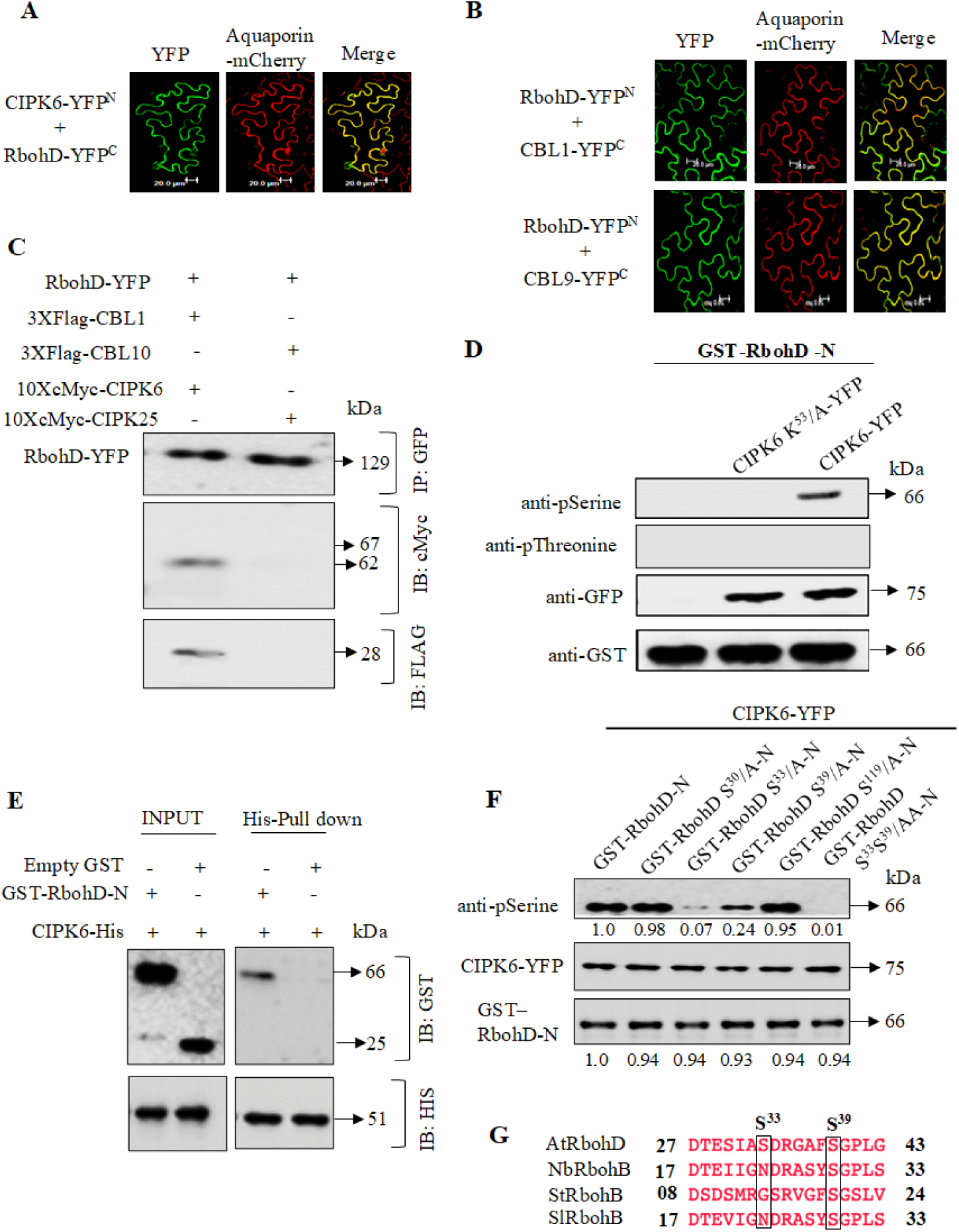
CBL1/9-CIPK6 module interacts with RbohD *in vivo* and CIPK6 phosphorylates the N -terminus of RbohD. **(A)** CIPK6-YFP^N^ interacts with RbohD-YFP^C^ at PM in *N. benthamiana* leaves by BiFC. Interaction at PM revealed by co-localization with PM marker Aquaporin-mCherry. Reconstitution of truncated YFP (green) merged with PM marker Aquaporin-mCherry (red) produced yellowish appearance. **(B)** RbohD-YFP^N^ interacts with CBL1-YFP^C^ and CBL9-YFP^C^ at PM *in planta*. Truncated YFP showed reconstitution of YFP (green) and merged with PM marker Aquaporin-mCherry (red). **(C)** Co-IP showed that CIPK6 and CBL1 interacted with RbohD. Combination of 10xcMyc-CIPK6, 3xFlag-CBL1 and RbohD-YFP and 10xcMyc-CIPK25, 3xFlag-CBL10 and RbohD-YFP were transiently expressed in *N. benthamiana* leaves. **(D)** In an *in vitro* kinase assay, only the serine residues in the N-terminus of RbohD (RbohD-N) are phosphorylated by CIPK6 (Active). CIPK6 (Active) phosphorylated the serine residues in RbohD-N only whereas CIPK6 K^53^/A (Inactive) does not phosphorylate either the serine or the threonine residues in the RbohD-N. **(E)** *In vitro* pull-down assay shows recombinant GST-RbohD-N directly interacts with His-CIPK6. Immunoblots were developed with anti-GST and anti-His antibody. **(F)** Cipk6 phosphorylates serine residues present at 33 (S^33^) and 39 (S^39^) positions in RbohD-N. When S^33^ and S^39^ were substituted with alanine independently, anti-pSerine antibody developed through HRP-sec antibody showed faint signal. Phosphorylation signal was not observed, when S^33^ S^39^ both were substituted simultaneously with alanine. Intensity of Protein band showed below. **(G)** Multiple sequence alignment of AtRbohD, NtRbohB, StRbohB and SlRbohB revealed that S^33^ is unique in Arabidopsis RbohD whereas S^39^ is conserved in all aligned Rboh.

Phosphorylation of Ser^39^ of RbohD by BIK1 during PTI has been previously reported (Kadota et al., 2014; Li et al., 2014) while, Ser^33^ has not been reported so far as the target of any kinase. Additionally, sequence alignment of RbohB/D orthologs across multiple plant species revealed that Ser^39^ is conserved, whereas Ser^33^ is unique to Arabidopsis. This suggests a species-specific regulatory mechanism in ROS-mediated immune responses (Figure 5G).Our study provides new insights into the phosphorylation-mediated regulation of RbohD by CIPK6, highlighting Ser^33^ as a novel phosphorylation site. These findings contribute to the broader understanding of how CIPKs regulate ROS production in plant immunity.

### CBL1/9-CIPK6 mediated phosphorylation inhibits RbohD activity and negatively regulate host immunity

Biological significance of CIPK6-mediated phosphorylation of RbohD was investigated by trans-complementation assay in *N. benthamiana* leaves as described previously (Lee et al., 2020). Expression of *N. benthamiana* RbohB, the ortholog of *Arabidopsis* RbohD, was silenced by virus-induced gene silencing (VIGS). This was followed by transient expression of *Arabidopsis* RbohD in the silenced *N. benthamiana* plant. The trans-complemented plants were tested for ROS production after infiltration with *Pst* DC3000. Trans-complementation with *Arabidopsis* RbohD restored only the PTI-mediated H_2_O_2_ production in the RbohB-silenced lines. Co-expression of CIPK6 with RbohD led to less H_2_O_2_ production and co-expression of CIPK6 and CBL1 together with RbohD resulted in further lesser H_2_O_2_ production (Supplementary Figure 11A and 12A) upon infiltration with *Pst* DC3000, suggesting CIPK6 along with CBL1 negatively regulates RbohD activity during plant immune response. To investigate the functional significance of phosphorylation of Ser^33^ and Ser^39^ in the activity of RbohD during the immune response, these amino acids were replaced individually or in combination with alanine or aspartic acid to create phosphonull (S^33^/A; S^39^/A; S^33^S^39^/AA) or phosphomimetic (S^33^/D; S^39^/D; S^33^S^39^/DD) mutants of RbohD. Trans-complementation of the RbohB-silenced *N. benthamiana* leaves with RbohD S^33^/A generated significantly high level of H_2_O_2_, while complementation with RbohD S^33^/D led to more than 4-fold lesser H_2_O_2_ production demonstrating that phosphorylation of RbohD Ser^33^ by CBL1-CIPK6 negatively regulates PTI-mediated RbohD activity during immune response. Contrastingly, trans-complementation with RbohD S^39^/A generated two-fold lower amount of H_2_O_2_ while, complementation with RbohD S^39^/D showed significantly high level of H_2_O_2_ production (Supplementary Figure 11B and 12 B). BIK1-mediated phosphorylation of RbohD Ser^39^ was reported to be required for PAMP-induced ROS burst (Kadota et al., 2014; Li et al., 2014). Our result also supports that phosphorylation of RbohD Ser^39^ is necessary for *Pst* DC3000-induced PTI-mediated H_2_O_2_ production. We then trans-complemented the RbohB-silenced *N. benthamiana* leaves with the double-alanine or double-aspartate substituted RbohD (S^33^S^39^/AA; S^33^S^39^/DD) and infiltrated with *Pst* DC3000. To our surprise, complementation of *N. benthamiana* leaves with the double-alanine substituted RbohD (S^33^S^39^/AA) resulted in significantly high PTI-mediated production of H_2_O_2_ while, a compromised PTI-mediated H_2_O_2_ production was observed in case of the phosphomimetic mutant (S^33^S^39^/DD) of RbohD similar to the level of the single mutant RbohD S^33^/D (Supplementary Figure 11C and 12C). The same experiments were repeated using the *Arabidopsis* T3 homozygous lines expressing the wild type and the mutated versions of RbohD under *rbohD* mutant background. Similar results were obtained from the *Arabidopsis rbohD* mutant lines stably expressing *RbohD*, *RbohD*(*S^33^A*), *RbohD(S^39^A)*, *RbohD*(*S^33^S^39^/AA*), *RbohD*(*S^33^D*), *RbohD*(*S^39^D*), *RbohD*(*S^33^S^39^/DD*) (Figure 6, A-D and Supplementary Figure 13, A-F). Taken together, this result demonstrated that CIPK6-mediated phosphorylation of Ser^33^ negatively regulated RbohD-mediated ROS burst. Further, negative regulation of RbohD activity due to CIPK6-mediated Ser^33^ phosphorylation superseded the Ser^39^ phosphorylation-mediated activation of RbohD. Overall, these findings reveal that CIPK6-mediated phosphorylation of RbohD serves as a crucial checkpoint in plant immunity, ensuring that ROS production is tightly controlled process.

**Figure 6:**
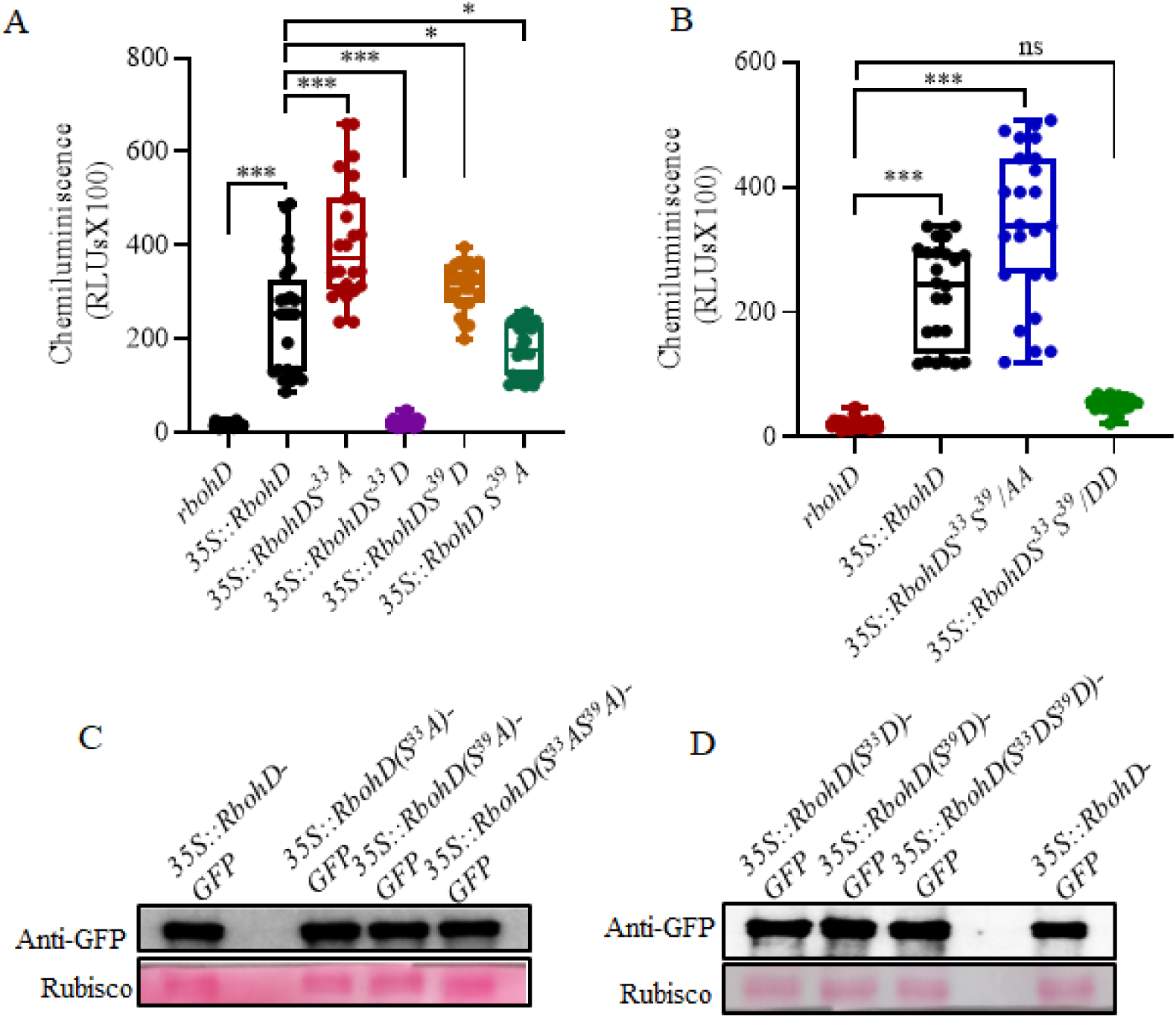
CIPK6 negatively regulates ROS generation via RbohD in Arabidopsis. (A) After *Pst*DC3000 challenge, theosingle phospho-null mutant *35S::AtRbohD S^33^/A* exhibited higher ROS production compared to *35S::AtRbohD S^39^/A* and *35S::AtRbohD*, whereas the single phospho-mimetic mutant *35S::AtRbohD S^33^/D* completely abolished ROS generation in comparison to *35S::AtRbohD S^39^/D* and *35S::AtRbohD.* The box plot illustrates the maximum ROS levels reached during PTI. An immunoblot using anti-GFP antibodies confirms the equal expression of AtRbohD, AtRbohD S^33^/A, AtRbohD S^39^/A, AtRbohD S^33^/D, and AtRbohD S^39^/D. (B) The over expression of double phospho-null mutant *35S::AtRbohD S^33^S^39^/AA* exhibited higher ROS production compared to *AtRbohD* over expressing plants, whereas the overexpression of double phospho-mimetic mutant *35S::AtRbohD S^33^S^39^/DD* displayed reduced ROS production following *Pst* inoculation. The box plot shows the maximum ROS achieved during PTI. (C) and (D) An immunoblot using anti-GFP antibodies confirms the equal expression of AtRbohD, AtRbohD S^33^/A, AtRbohD S^39^/A, AtRbohD S^33^/D, and AtRbohD S^39^/D, *AtRbohD S^33^S^39^/AA, and AtRbohD S^33^S^39^/DD*.

## Discussion

Apoplast-associated ROS production by plasma membrane NADPH oxidases, namely respiratory burst oxidases (Rbohs) is strongly associated with plant defence response. In this regard, RbohD and RbohF are the most studied Rboh family members. Deletion of functional RbohD eliminates majority of the pathogen-triggered ROS production (Torres et al., 2002).

Calcium (Ca^2+^) signalling plays a crucial role in RbohD activation (Dubiella et al., 2013; Li et al., 2014). Ca^2+^ binding to the N-terminal EF hand of RbohD induced conformational changes in the protein (Ogasawara et al., 2008). In addition, RbohD has several potential phosphorylation sites which are targeted by various kinases (Dubiella et al., 2013; Li et al., 2014; Kadota et al., 2014; Lee et al., 2020; Kimura et al., 2020; Zhang et al., 2018; Chen et al., 2017) indicating multiple layers of regulation of its activity. Plant innate immunity is a precisely controlled process of positive and negative regulation. Absence of negative regulation would cause over- and continuous activation of defence response resulting in retarded growth and even death to the plants (Heil and Baldwin, 2002; Nimchuk et al., 2015). Negative regulation of RbohD activity due to phosphorylation at its C-terminus by PBL13 and its subsequent ubiquitin-mediated degradation has been reported recently (Lee et al., 2020). Our study reports a direct and *in planta* interaction of CIPK6 and CBL1 or -9 with the plasma membrane-localized RbohD. This interaction led to the negative regulation of RbohD activity due to CIPK6-mediated phosphorylation of Serine^33^ at its N-terminal cytoplasmic domain. CIPK6 was also able to phosphorylate Serine^39^ of RbohD leading to its activation. Serine^39^ of RbohD has been reported as the target of PAMP-dependent phosphorylation and activation by BIK1 kinase, a component of pattern recognition receptor (PRR) complex (Kadota et al., 2014; Li et al., 2014). Our result also supports activation of RbohD by Serine^39^ phosphorylation. However, we show that Serine^33^ is the major CIPK6-mediated phosphorylation site and CIPK6-mediated Serine^33^ phosphorylation and inactivation of RbohD supersedes its activation due to Serine^39^ phosphorylation. *Pst* infection quickly downregulated CIPK6 mRNA and protein abundance. Reduction in CIPK6 activity would lead to reduction in Serine^33^ phosphorylation. Subsequently, PAMP-triggered BIK1-mediated phosphorylation of Serine^39^ and the other residues of RbohD (Kadota et al., 2014; Li et al., 2014) would lead to the activation of RbohD in addition to the phosphorylation and activation by other kinases (Dubiella et al., 2013; Ogasawara et al., 2008; Kobayashi et al., 2007). For example, bacterial infection elicits release of ATP and ATP-activating kinase DORN1 activates RbohD by phosphorylating its Ser^22^ and Thr^24^ to induce stomata closure (Chen et al., 2017). Different mechanisms of Rboh activation may exist in other species. Such as, unlike downregulation of CIPK6 expression in *Arabidopsis*, SlCIPK6 expression was quickly upregulated by *Pst* infection and SlCIPK6 positively regulated ROS production in tomato (de la Torre et al., 2013). Unlike Serine^33^, Serine^39^ is a conserved amino acid among the functional RbohD orthologues across the species. A preliminary search identified a Serine corresponding to Serine^33^ only in a RbohD ortholog in *Brassica napus* (NCBI: XP_013644255.1), which belongs to Brassicaceae family like *Arabidopsis*. Unlike CBL10 of tomato, *Arabidopsis* CBL10 did not interact with CIPK6 and the *cbl10* mutant plant did not show any resistance to *Pst* infection. Arabidopsis CBL10 has an unusually long and unique hydrophobic N-terminal domain (74 amino acid) (Weinl and Kudla, 2009). Arabidopsis CBL10 protein localizes to vacuolar membranous invaginations surrounding the nucleus and mobile punctate structures in the cytoplasm (Kim et al., 2007). N-terminal domain of tomato CBL10 shows a very low level of sequence similarity to its Arabidopsis ortholog and is localized to plasma membrane (de la Torre et al., 2013). Therefore, it appears that pathogen-triggered immune response in plants varies with the species.

Arabidopsis CBL1 and CBL9 were shown to interact with CIPK26 and CIPK26-CBL1/9 complex was shown to phosphorylate RbohF *in vitro* and activate RbohF in a heterologous system human embryonic kidney cell line (HEK29) (Drerup et al., 2013) CBL1-CIPK26-mediated phosphorylation also enhanced RbohC activity in HEK29 cell line (Zhang et al., 2018). Tomato CBL10 and CIPK6 act as positive regulators of ROS production and *Pst* DC3000-triggered programmed cell death (de la Torre et al., 2013). Our result showed mutated Arabidopsis line with disrupted CBL10 produced similar bacterial growth as Col-0 upon *Pst* DC3000 infiltration. There was, however, no report so far on involvement of CBL1 and its paralog CBL9 in plant immune response. In this manuscript, we showed CBL1 and CBL9 redundantly negatively regulate plant immune response and immunity in Arabidopsis. Expression of any of these two protein molecules in the double mutant (*cbl1cbl9*) restored the wild type phenotype. Arabidopsis do not induce ETI-triggered immunity against AvrPto. Infiltration with *Pst* DC3000 harbouring an avirulent effector AvrRPS4 showed enhanced ETI-triggered response in the double mutant suggesting a role of CBL1/ 9 in regulating both PTI- and ETI-mediated immune response in Arabidopsis.

CBL1/9, CIPK6 and RbohD were identified in a membrane-bound complex. CIPK6 itself is not a membrane bound protein and it requires a membrane-localised CBL to localize close to plasma membrane to form a complex (Held et al., 2011). This was shown by co-expressing CIPK6 and CBL1/9 having a myristoylation-target glycine residue replaced with alanine. CBL1 and CBL9 could interact with CIPK6 irrespective of their subcellular localization as shown by the interaction of their G^2^/A mutants with CIPK6 in nucleus and cytoplasm. Our results showed the kinase activity of CIPK6 was required for the negative regulation of plant immunity and ROS production. CIPK6 T^182^/DΔC lacks the NAF domain required for CBL binding. In spite of that, expression of this highly active version of CIPK6 was able to restore the bacterial growth in the *cipk6* mutant and showed low ROS production. This is probably due to high cytoplasmic kinase activity of this CIPK6 variant that was able to phosphorylate and inhibit the cytoplasmic domain of RbohD. In a natural situation, plasma membrane localization and close proximity of relatively low active wild type CIPK6 is required for phosphorylation-mediated inhibition of RbohD. The extent of ROS production and defence response varied significantly with the relative levels of kinase activities of the CIPK6 variants (WT, T^182^/D, K^53^/A and T^182^/DΔC). However, in spite of very high kinase activities of CIPK6 T^182^/D and CIPK6 T^182^/DΔC the ROS activity did not decrease and bacterial growth did not increase significantly in these plants over the *cipk6*/CIPK6 plants indicating a saturation in the negative regulation of RbohD activity and immunity by CIPK6.

Influx of Ca^2+^ across the plasma membrane is essential for PAMP-driven ROS burst and Calcium-dependent signalling components are required for the rapid signal propagation and activation of the immune response mechanisms (Boudsocq et al., 2010; Dubiella et al., 2013). CBL-family of proteins are Calcium-sensors and Calcium-binding proteins, and CBL-CIPK modules requires Ca^2+^ for their *in vitro* kinase activity (Guo et al., 2001; Luan, 2009; Chaves-Sanjuan et al., 2014). Our result report a direct involvement of a Calcium-sensor protein and its interacting protein kinase in negatively regulating of ROS production by phosphorylating and inactivating RbohD in Arabidopsis. CBL and CIPK family of genes were mostly studied in relation to various abiotic stresses (Liu et al., 2000; Qiu et al., 2002; Batelli et al., 2007; Li et al., 2014; Tang et al., 2020). Comparatively, reports on their involvement in pathogen-triggered immune response is scars. We showed that CBL1 and 9, like their interacting kinase CIPK6, are the negative regulators of immunity in Arabidopsis. Many membrane-localised transporters were identified as the substrates of CBL-CIPK modules and in most of the cases interaction or CIPK-mediated phosphorylation activated the substrates (for review, Tang et al., 2020). Our study identified a new substrate of Arabidopsis CIPK6 and a direct role of CBL1/9-CIPK6 kinase module in regulating immune response in *Arabidopsis thaliana* (Supplementary Figure 14).

This work advances our understanding of how Ca^2+-^dependent signaling pathways fine-tune immune responses by balancing activation and inhibition of ROS production. While CBL-CIPK modules have been extensively studied in abiotic stress responses, their role in plant immunity remains largely unexplored. Our findings introduce CIPK6 as a novel negative regulator of plant defense and expand the functional repertoire of CBL1 and CBL9 in immune signaling. By identifying a species-dependent regulation of CIPK6 and its interaction with CBL proteins, this study also highlights the evolutionary divergence of immune signaling pathways in different plant species. These insights could aid in developing strategies to manipulate ROS-mediated defense responses for improved disease resistance in crops.

## Materials and Methods

### Plant materials, growth conditions and transformation

Arabidopsis plants were grown in agropeat soil under controlled growth chamber (Conviron, Winnipeg, Canada) at 21-23 ^0^C temperature, 70% humidity, 10 h light /14 h dark photoperiod and 100 µmol µm^-2^ S^-1^. *N. benthamiana* plants were grown in garden soil under controlled growth condition at 25 ^0^C temperature, 50% relative humidity, 10 h light /14 h dark photoperiod and 100 µmol µm^-2^ S^-1^. All *Arabidopsis* genotypes were of Col-0 ecotype. T-DNA insertion lines *cipk6* (GK-448C12-CS442948), *cbl1* (SALK_110426), *cbl9* (SALK_142774) and *cbl10* (SALK_056042) have been described earlier (Sardar et al., 2017; Held et al., 2011; Ren et al., 2013; Xu et al., 2006). The mutant *rbohD* (CS_68522) was procured from Arabidopsis Biological Resource Center (ABRC), Columbus, OH. Mutants *cbl1*, *cbl9*, *cbl10* were procured from Prof. Wei-Hua Wu, China Agricultural University, China. CIPK6, CBL1 and CBL9 and RbohD variants (CIPK6 T^182^/D, CIPK6 K^53^/A, CBL1 G^2^/A, CBL9 G^2^/A, RbohD S^33^/A, RbohD S^33^S^39^/AA, RbohD S^33^/D, RbohD S^33^S^39^/DD) were created by the site directed mutagenesis as previously described by Meena et al., 2015. All the primers used for this purpose has been described in the supplementary table 1. CIPK6, CBL1, CBL9, RbohD and their variants were expressed under 35CaMVS promoter using the plasmid pGWB2 in the plants. *Agrobacterium tumefaciens* strain GV3101 harbouring *CIPK6, CBLs and RbohD* constructs were introduced in the *cipk6, cbl1cbl9* and *rbohD* mutant plants, respectively, by floral dip as described previously (Clough and Bent, 1998). The transformed seeds were selected on half-MS/kanamycin plate and the presence of *CIPK6*, CBL, RbohD DNA fragments in the transformed plant lines was confirmed by genomic DNA PCR. Expression was confirmed by Semi-quantitative RT-PCR. T3 single-insertion homozygous seeds were used for all *Arabidopsis* experiments.

### RNA extraction and q-PCR

Total RNA was isolated from Arabidopsis leaves using the Trizol method (Sigma), and any DNA contamination was removed through DNase treatment thorugh DNAse enzyme (Ambion) at 25^0^C for 20 minutes followed by stopping the reaction by 0.5mM EDTA at 72^0^C before cDNA synthesis. Then cDNA was synthesized from 1-3 µg of RNA using the Thermo Verso cDNA synthesis kit. Quantitative real time PCR (qPCR) was performed using 50-100 ng of cDNA, Power SYBR Green Master Mix (Applied Biosystems, USA), and an ABI-PRISM 7500 FAST sequence detector. Each qPCR analysis included three biological replicates, with two technical replicates for each cDNA sample. ACTIN2 was used as house-keeping gene.The primers used for expression analysis are listed in Supplementary Table S1.

### Subcellular localization and Bimolecular Fluorescence Complementation (BiFC)

*CBL1*, *CBL9*, *CBL1 G^2^/A* and *CBL9 G^2^/A* coding sequence were cloned in pEG101 by gateway technology (Invitrogen, Waltham, MA). Aquaporin-mCherry was used as plasma membrane marker and NLS-RFP (Nuclear localization signal-Red Fluorescence Protein) was used as nucleus marker (Nelson et al., 2007). *Agrobacterium* strain EHA105 carrying *CBL* constructs, marker constructs were mixed at 1:1 v/v ratio and incubated for 2-4 hours at 23 ^0^C in infiltration media (10 mM MgSO_4_, 10 mM MES 100 µM Acetosyringone). The mix was infiltrated in *N. benthamiana* leaves and kept in growth-room for 3 days at 25 ^0^C. The expression was monitored by the fluorescence captured by a confocal laser scanning microscopy Leica TCS SP5 (Leica microsystems, Wetzlar, Germany) with excitation wavelength of 514 nm for YFP and 543 nm for mCherry and RFP.

For the Bimolecular fluorescence complementation (BiFC), *CIPK6* and *CIPK6 K^53^/A* coding sequences were cloned into pSITE-nEYFP-C1 (N-terminal YFP fragment) and *CBL1*, *CBL9*, *CBL1 G^2^/A*, *CBL9 G^2^/A* and *RbohD* coding sequences were cloned in pSITE-cEYFP-N1 (C-terminal YFP fragment) by the gateway technology. *RbohD* was also cloned in pSITE-cEYFP-N1 (C-terminal YFP fragment). *Agrobacterium* strain EHA105 carrying combination of *CBLs-YFP^C^* + *CIPK6-YFP^N^*, *CIPK6-YFP^N^* or *CIPK6 K^53^/A-YFP^N^* + *RbohD-YFP^C^*, *CBLs-YFP^C^* + *RbohD-YFP^N^* constructs. The constructs were infiltrated in *N. benthamiana* leaves and reconstitution of fluorescence was monitored as described above. Subcellular fractionation of CBLs expressing *N. benthamiana* leaves was performed using Qproteome Cell Compartment Kit following protocol as mentioned in the manufacturer’s instructions manual (Qiagen, GmbH, Germany). The primers are enlisted in the supplemenatry table S1.).

### Bacterial infection and SA Quantification

Bacterial inoculation, determination of growth and quantification of salicylic acid was performed as described by Sardar et al., 2017. Briefly, the bacterial pathogen *Pseudomonas syringae* pv. *tomato* DC3000 (*Pst* DC3000) alone or carrying a construct encoding *AvrRps4* were grown on King’s medium B agar plates or in liquid medium supplemented with 50 µg/ml rifampicin and 50 µg/ml kanamycin at 28 ^0^C. The bacterial culture was resuspended in 10 mM MgCl_2_ and manually infiltrated in leaves with an OD_600_ of 0.0005 for the virulent strain, and OD_600_ of 0.001 for the avirulent strain. Crushed leaf samples were serially diluted with 10 mM MgCl_2_ and plated onto King’s B medium containing the appropriate antibiotics for bacterial counts. The values presented are the mean of at least three biological replicates. At least 8 representative leaves for each replicate and five technical replicates were used to generate results. Statistical analyses were performed using Two-way ANOVA (Fujikoshi Y, 1993).

### Yeast two-hybrid (Y2H) assay

Y2H was performed as described by (Meena et al., 2019). Briefly, the full-length coding sequences of CIPK6 and all CBLs (CBL1–CBL10) were fused to the GAL4 DNA-binding domain (BD) and activation domain (AD), respectively in the yeast Y2H plasmids pGBKT7-BD and pGADT7-AD, respectively (Takara Bio Inc., Kusatsu, Japan). The combinations of constructs were co-introduced into the Y2H Gold strain of yeast using the Matchmaker Gold yeast Two-Hybrid system kit (Takara Bio Inc.). The yeast colonies harboring both the plasmids were selected on double drop-out (DDO) medium lacking leucine and tryptophan and were confirmed by colony PCR. Positive colonies selected on DDO plates were grown in liquid DDO media for 2 days, adjusted to an OD_600_ of 1, 0.1, and 0.01. A 10 μl aliquot of each dilution was spotted on quadruple drop-out medium (-Leu-Trp-Ade-His) with 2.5 mM 3-AT (3-amino-1,2,4-triazole). Plates were incubated at 30 ^0^C for 5 days.

### Co-immunoprecipitation, *in vitro* pull-down and western blot

To investigate interactions between CIPK6, CBL1, CBL9 and RbohD, *Agrobacterium* (EHA105) cultures carrying plasmid constructs encoding *10xcMyc-*fused *CBL1/9* and *CIPK6* fused to *YFP* or *CIPK25* fused to *YFP* were co-infiltrated in *N. benthamiana* leaves. To detect the *in vivo* interaction of CIPK6 and CBL1/9 with RbohD, *Agrobacterium* (EHA105) carrying combination of *10xcMyc-CIPK6*, *3xFlag-CBL1* and *RbohD-YFP* and combination of *10xcMyc-CIPK25*, *3xFlag-CBL10* and *RbohD-YFP* were infiltrated in *N. benthamiana* leaves. After 3 days, leaves sample were harvested and protein were isolated in 1 ml/ gm plant protein extraction buffer (100 mM Tris-HCl, pH 7.5, 150 mM NaCl, 5 mM EDTA, 10% glycerol, 0.1% Triton X-100, 1x plant protease inhibitors, and 1 mM phenylmethylsulfonyl fluoride). 1 mg of supernatant crude protein was incubated with 1µl GFP antibody (Abcam, Cambridge, UK) and Sepharose beads (GE Healthcare, Chicago, USA) for 8 h at 4 ^0^C. Sepharose beads bound proteins were washed with washing buffer (1x PBS, 1x protease inhibitor and 0 .1% tween 20) to remove nonspecific bound protein. Proteins bound with beads were boiled in 50 µl of 1x laemmli sample buffer and detected by immunoblotting. Primers attached in the supplementary table S1.

For *in vitro* pull-down assays, pGEX4T2 without or with RBOHD-N for GST (Glutathione-S-transferase)-fused protein expression, and pET28a with CIPK6 for 6xHis (Histidine)-fused protein expression were expressed in *E. coli*. Supernatant fraction of the bacterial lysate containing His-CIPK6 was incubated with Ni-NTA beads. Affinity purified GST or GST-RbohD-N were incubated with 6xHis-CIPK6 immobilized on Ni-NTA beads (Qiagen) for 3 hrs. The beads were washed with washing buffer (1x PBS, 0.01% tween 20). Proteins bound with beads were detected by immunoblotting with anti-GST antibody (Sigma-Aldrich, St. Louis, MI) and anti-His antibody (Sigma-Aldrich, St. Louis, MI).

### Kinase assay

For CIPK6 auto-kinase assay, purified bacterially expressed proteins were used for *in vitro* kinase assays. Coding sequences of *CIPK6* and variants (*CIPK6* K^53^/A, *CIPK6* T^182^/D, *CIPK6*ΔC, *CIPK6* T^182^/DΔC), *RbohD-*N (1-376 aa) and *CBL1/9* were cloned in pGEX4T2 bacterial expression vector and introduced into *Escherichia coli* BL21-CodonPlus (Agilent Technologies, Santa Clara, CA). Protein expression was induced using 1 mM IPTG (Isopropyl-β-D-thiogalactopyranoside). The protein was affinity purified using Glutathione-Sepharose beads (Bio-Vision, Waltham, MA) *via* eluting in 50 mM Tris-HCL pH-9.5, 3 mg/ml reduced Glutathione containing elution buffer. 6xHis-tagged CIPK6 was purified through Ni-NTA (Nickel-Nitrilotriacetic acid) beads (Quigen, Germany) and eluted by using elution buffer (50 mM Tris -HCl pH 7.4, 200 mM NaCl, 250 mM Imidazole). Primers are enlisted in the supplementary table S1.

For auto-phosphorylation activity assay, CIPK6 was used as both the enzyme and substrate. Kinase reaction was set in kinase buffer (20 mM Tris-Cl, pH 8.0, 5 mM MnCl_2,_ 1 mM CaCl_2_, 0.1 mM EDTA, 1 mM DTT, and 10 µCi γ-^32^P ATP) in 20 µl final volume and incubated at 30 ^0^C for 30 minutes. The amount of kinase and substrate were used in 1:10 ratio. Kinase reactions were terminated by adding 1x Laemmli sample buffer and was analysed by 12% SDS-PAGE. For the transphosphorylation assay, CIPK6 and its variants were expressed as YFP-fused protein in *N. benthamiana* leaves and immunoprecipitated with anti-GFP antibody and used as kinase. Myelin basic protein (MBP) or GST-RbohD-N was used as the substrate. 1 mg plant protein extract was used for immunoprecipitation. The washed protein-bound bead pellet was used in the kinase reaction. Radiolabelled substrates were detected and measured by autoradiography using Typhoon FLA9500 phosphoimager (GE Healthcare, Uppsala, Sweden) for the kinase activity of CIPK6. For the non-radiolabelled reactions, phosphorylated products were detected by immunoblotting with anti-pSerine and anti-pThreonine antibodies (Santa Cruz, Dallas, TX).

To study the CIPK6 enzyme kinetics in planta, *Agrobacterium* (EHA105) carrying 10xcMyc-CIPK6 was agro-infiltrated in *cipk6* plants leaves. After 60 h of infiltration, *Pst* DC3000 was infiltered in same leaves. Samples were harvested at 0,15, 45-, 90-, and 180-minutes post infiltration. Protein extracts were used either for immunoblot for detecting CIPK expression or CIPK6 was immunoprecipitated with anti-cMyc antibody (Sigma-Aldrich) for kinase activity.

### Mass spectrometry

To identify CIPK6-mediated phosphorylation site/s in RbohD-N, the kinase reactions were electrophoresed by 10% SDS-PAGE and stained with Coomassie brilliant blue. Protein band was excised and in-gel digestion was performed using trypsin (Promega, Madison, WI) following reduction with 5mM TCEP (Tris(2-carboxyethyl) phosphine) and alkylation with 50 mM iodoacetamide. Digested protein samples were cleaned using a C18 silica cartridge to remove the salt and dried using a speed vac. The dried pellet was resuspended in buffer A (5% acetonitrile, 0.1% formic acid). Mass spectrometry was performed using EASY-nLC 1000 system (Thermo Fisher Scientific, Waltham, MA) coupled to Thermo Scientific Q Exactive equipped with nanoelectrospray ion source. 1.0 µg of the peptide mixture was resolved using 50 cm PicoFrit column filled with 3.0 µm C18-resin. The peptides were loaded with buffer A (5% acetonitrile, 0.1% formic acid) and eluted with a 0–40% gradient of buffer B (95% acetonitrile, 0.1% formic acid). MS data was acquired using a data-dependent top10 method dynamically choosing the most abundant precursor ions from the survey scan. MS data were analysed with Proteome Discoverer (v2.2) against the *reference* Arabidopsis customized sequence database.

### VIGS and ROS burst measurement

The 400 bp non-conserved fragments of *NbRbohB* was PCR amplified from the total cDNA of *N. benthamiana* and sub-cloned into pTRV2 vector. To silence *NbRbohB* by virus induced gene silencing (VIGS), *Agrobacterium* (EHA105) culture carrying *pTRV1* and *pTRV2: NbRbohB* (OD_600_ = 0.8 -1.0) was agroinfiltrated in the leaves of 2 weeks old *N. benthamiana* seedling. After three weeks of agroinfiltration, newly emerged leaves were checked for *NbRbohB* silencing by RT-PCR. *Agrobacterium* (EHA105) carrying binary plasmid constructs with the coding sequences of *CIPK6-YFP*, *CBL1-YFP*, *RbohD-YFP* and phospho null/mimic mutants of RbohD (*RbohD S^33^/A*, *RbohD S^33^S^39^/AA*, *RbohD S^33^/D*, *RbohD S^33^S^39^/DD*) were agroinfiltrated in *NbRbohB*-silenced *N. benthamiana* leaves or *rbohD* mutant of *Arabidopsis* in various combination, to detect ROS production. ROS measurement was performed in the stable lines of *cipk6/CIPK6*, *cbl1cbl9/CBL1/9* and their mutant variants. ROS detection and measurement was performed as described by Sardar et al., 2017. Briefly, leaf discs were excised with a cork borer (1.1 cm^2^) and floated adaxial side up overnight on sterile water. Prior to elicitation, water was replaced with 100 µl of *Pst*DC3000 (OD600= 0.1) and leaf discs were vacuum inoculated, further 100 µl of the elicitation solution containing 0.2 μM luminol (Sigma-Aldrich), 20 μg ml^−1^ horseradish peroxidase (HRP; Sigma-Aldrich was put in each well of 96 well flat bottom Luminescence plate. *Pst*-induced *in vivo* ROS production was measured as relative luminescence unit per sec (RLU/sec) using a POLAR star Omega (BMG Labtech, York, UK) 96-well microplate luminometer. The expression of CIPK6-YFP, CBL1-YFP and RbohD-YFP proteins in the leaf discs was detected by immunoblotting using anti-GFP antibody (Abcam, Cambridge UK).

## Supplementary Data

**Figure S1.** Interaction of CIPK6 with various CBLs (1-10).

**Figure S2.** Subcellular fractionation of CBL1-YFP and CBL9-YFP.

**Figure S3.** CBL1 and CBL9 are the negative regulators of immunity in *Arabidopsis thaliana* against Pst DC3000 infection.

**Figure S4.** CBL1 and CBL9 are the negative regulators of immunity in *Arabidopsis thaliana* against *Pst* DC3000 (AvrRPS4) infection.

**Figure S5.** CBL1 and CBL9 are the negative regulators of ROS production and salicylic acid in response to avirulent *Pst* DC3000 (AvrRPS4) infection.

**Figure S6.** Localization and subcellular fractionation of CBL9-YFP and CBL9 G^2^/A-YFP.

**Figure S7.** Interaction of CBL9 G^2^/A with CIPK6 by BiFC assay.

**Figure S8.** Altered localization of CBL1/9-CIPK6 complex affects its role in immune response.

**Figure S9.** Interaction of CIPK6 K^53^/A with RbohD by BiFC assay.

**Figure S10.** Mass spectrometry analysis of phosphorylated RbohD.

**Figure S11.** Time-course of luminol enhanced chemiluminescence assay of in vivo ROS production in *N*. *benthamiana TRV-NbRbohB* silenced plants.

**Figure S12.** CIPK6 and its interacting CBLs negatively regulates the ROS producing activity of RbohD in *Nicotiana*.

**Figure S13.** (A)-(C) Time course of ROS production in *Arabidopsis* plants, over expressing *RbohD*, *RbohD*(*S^33^A*), *RbohD(S^39^A)*, *RbohD*(*S^33^S^39^/AA*), *RbohD*(*S^33^D*), *RbohD*(*S^39^D*), *RbohD*(*S^33^S^39^/DD*), and *rbohD* mutants. (B) and (D) DAB staining. (E) Time course of ROS generation was done in *rbohD* mutants, *35S::RbohD-GFP*, CIPK6 overexpressing plants in the background of 35S::RbohD and CIPK6 and CBL1 overexpressing in the 35S::RbohD background. (F) Box plot to show the maximum ROS values in the indicated genotypes.

**Figure 14.** Proposed cartoon showing the regulation of RbohD-mediated ROS generation.

**Table S1.** List of primers used in this study

## Acknowledgements

The assistance of central instrumentation facilities, metabolomics facility and DISC of NIPGR in various experiments is acknowledged. The authors are grateful to the DBT-eLibrary Consortium (DeLCON) for providing access to e-Resources.

## Data availability

All data supporting the findings of this study are available within the paper and within its supplementary materials published online. The gene sequences are available in National Centre for Biotechnology Information (NCBI) (http://www.ncbi.nlm.nih.gov).

