## Supplementary file for "CBL1/9-CIPK6 complex negatively regulates Respiratory burst oxidase homolog D in *Arabidopsis thaliana*"

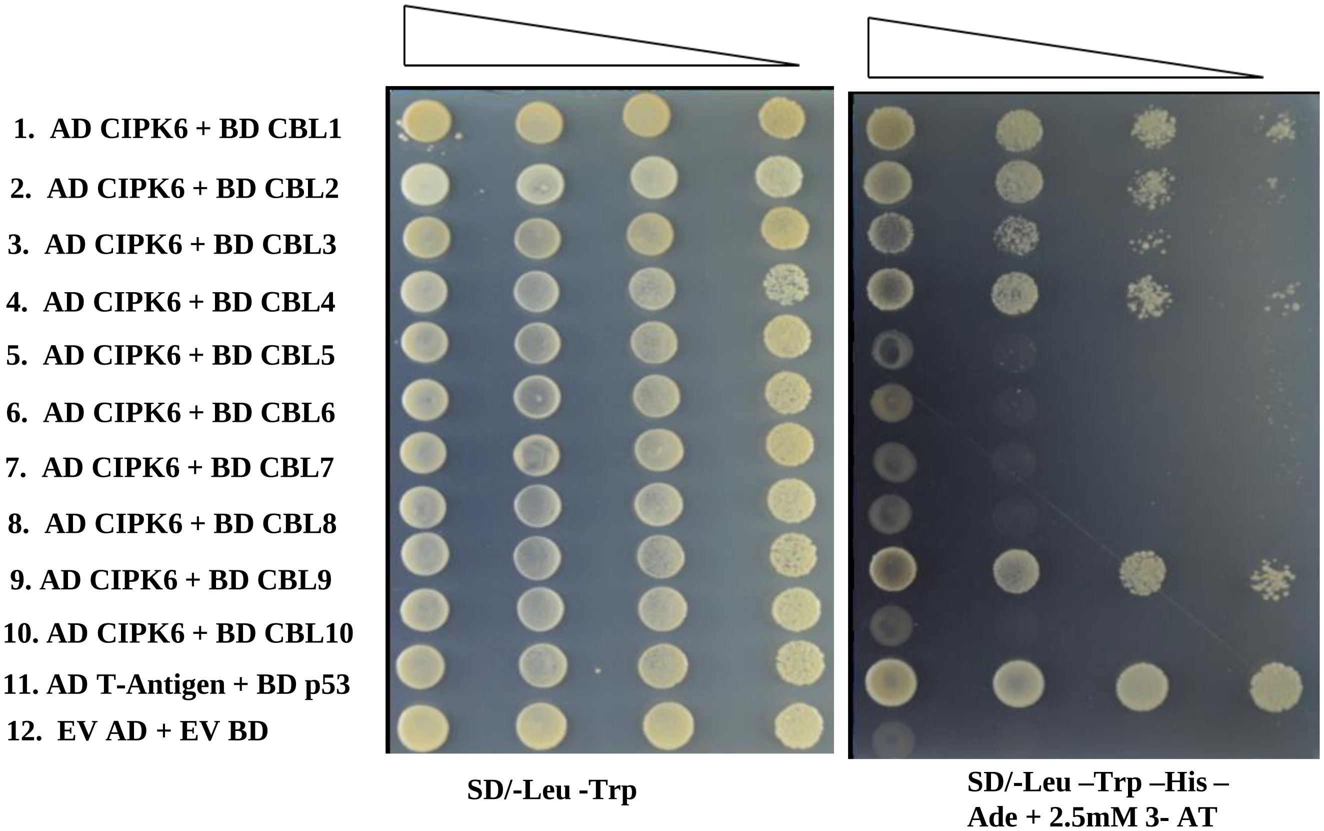

**Figure S1. Interaction of CIPK6 with various CBLs (1-10).** Yeast two hybrid assay shows interactions between CIPK6 and CBL proteins.Y2H gold strain was co-transformed with *pGADT7-CIPK6* + *pGBKT7-CBL1-10*, *pGADT7 T-Antigen* + *pGBKT7-P53*, and Empty *pGADT7* + Empty *pGBKT7*. Their Log phase culture was serially diluted and spotted on auxotrophic selection media.

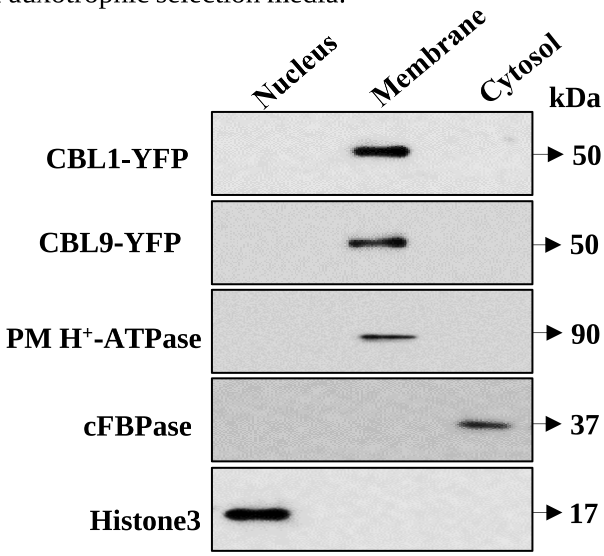

**Figure S2. Subcellular fractionation of CBL1-YFP and CBL9-YFP.** Immunoblot of subcellular fractions with anti-GFP antibody showing CBL1 and CBL9 are exclusively present in plasma membrane (PM). $H^+$-ATPase is used as a marker for the plasma membrane, Fructose 1,6-bisphosphatase (cFBPase) for the cytosol, and H3 histone for the nucleus. Here, subcellular fraction was probed using antibody against the above mentioned marker protein.

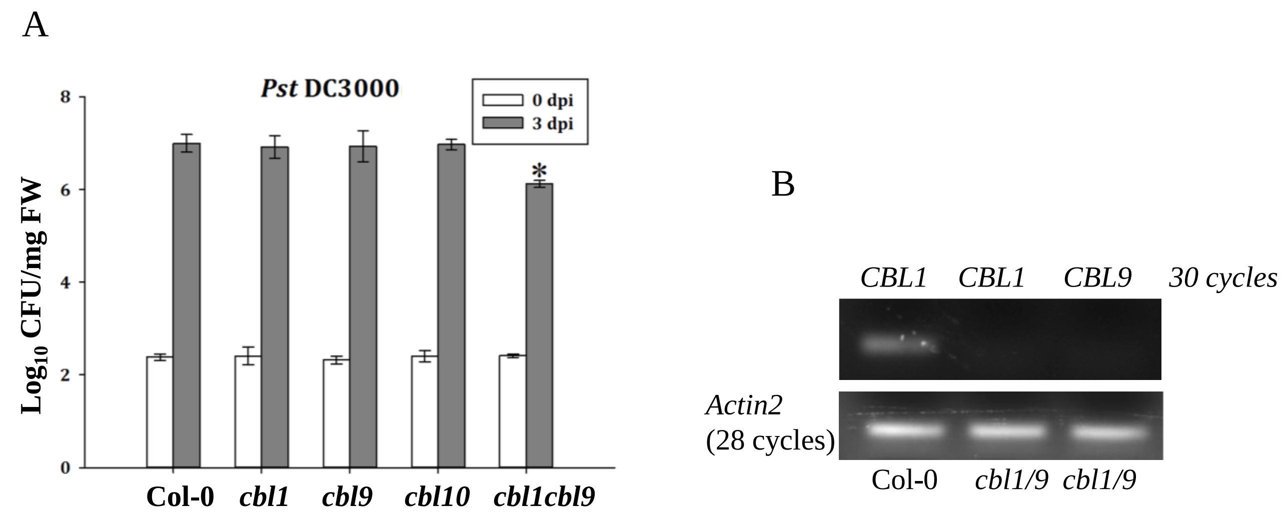

**Figure S3. CBL1 and CBL9 are the negative regulators of immunity in *Arabidopsis thaliana* against *Pst* DC3000 infection**. *Pst* DC3000 ($OD_{600}$ = 0.0005) was infiltrated in the leaves of *Arabidopsis* T-DNA insertion mutants *cbl1*, *cbl9*, *cbl10* and *cbl1cbl9*. Bacterial count (cfu/mg of fresh weight) was evaluated at 0 day and 3 days post infection (dpi) of. Asterisk shows significant difference following two-way ANOVA (α=0.05). (B) Semi q RT-PCR to check the *CBL1* and *CBL9* expression in *cbl1/9 d*ouble mutant plants.

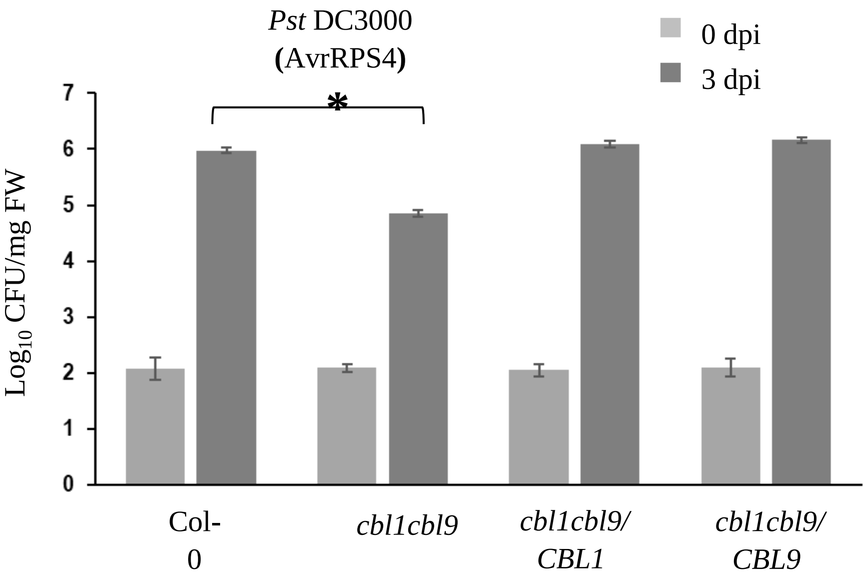

**Figure S4. CBL1 and CBL9 are the negative regulators of immunity in *Arabidopsis thaliana* against avirulent *Pst* DC3000 (AvrRPS4) infection. (A)** Bacterial count was evaluated in Col-0, *cbl1cbl9*, and these mutants complemented with CBL1 and CBL9 expression (*cbl1cbl9/CBL1*, and *cbl1cbl9/CBL9*) at 0 and 3 dpi after *Pst* DC3000 (AvrRPS4) ($OD_{600}$ = 0.001) infection. Asterisk shows significant difference following two-way ANOVA (α=0.05).

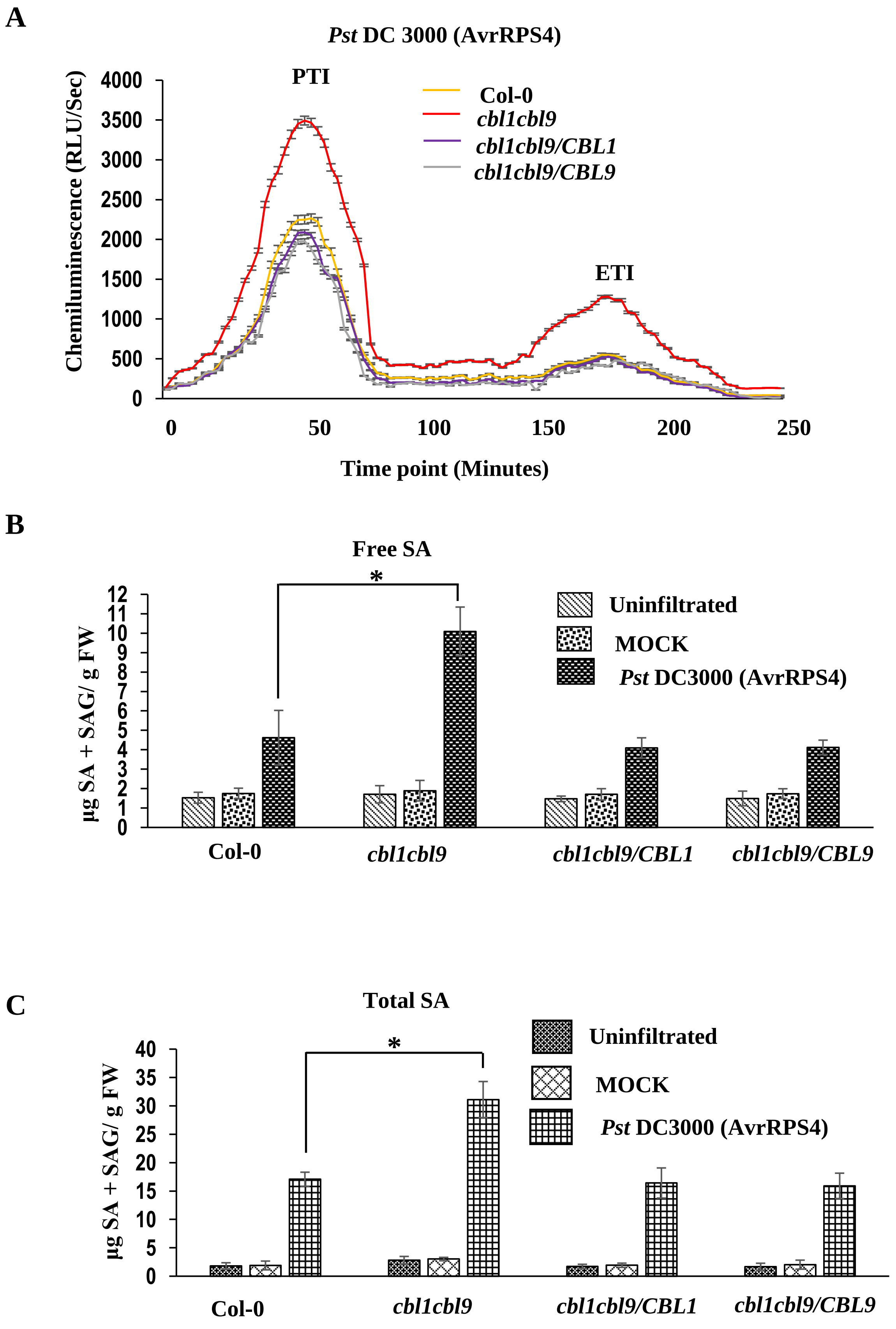

A
Pst DC 3000 (AvrRPS4)
PTI
ETI
Col-0
cbl1cbl9
cbl1cbl9/CBL1
cbl1cbl9/CBL9
Chemiluminescence (RLU/Sec)
4000
3500
3000
2500
2000
1500
1000
500
0
0
50
100
150
200
250
Time point (Minutes)
B
Free SA
*
Uninfiltrated
MOCK
Pst DC3000 (AvrRPS4)
µg SA + SAG/ g FW
12
11
10
9
8
7
6
5
4
3
2
1
0
Col-0
cbl1cbl9
cbl1cbl9/CBL1
cbl1cbl9/CBL9
C
Total SA
*
Uninfiltrated
MOCK
Pst DC3000 (AvrRPS4)
µg SA + SAG/ g FW
40
35
30
25
20
15
10
5
0
Col-0
cbl1cbl9
cbl1cbl9/CBL1
cbl1cbl9/CBL9

**Figure S5. CBL1 and CBL9 are the negative regulators of ROS production and salicylic acid in response to avirulent *Pst* DC3000 (AvrRPS4) infection**

**(A)** Time-course of *in vivo* luminol based ROS production assay in response to *Pst* DC3000 (AvrRPS4) ($OD_{600}$ = 0.02) inoculation in leaf discs of Col-0, *cbl1cbl9*, *cbl1cbl9/CBL1*, *cbl1cbl9/CBL9.* ROS was measured for 250 mins. Results are the mean (±SE), n>35 leaf disks. ROS peaks of PAMP-triggered and Effector-triggered immune (PTI, ETI) response were shown.

**(B)** Free salicylic acid (SA) and (C) total SA was measured in 4 week-old leaves of Col-0, *cbl1cbl9*, *cbl1cbl9/CBL1*, *cbl1cbl9/CBL9* plants at 24 hours post inoculation (hpi) of *Pst* DC3000 (AvrRPS4) ($OD_{600}$ = 0.001). 10 mM $MgCl_2$ was used as mock. The asterisks show a significant difference following the two-way ANOVA ($\alpha$=0.05).

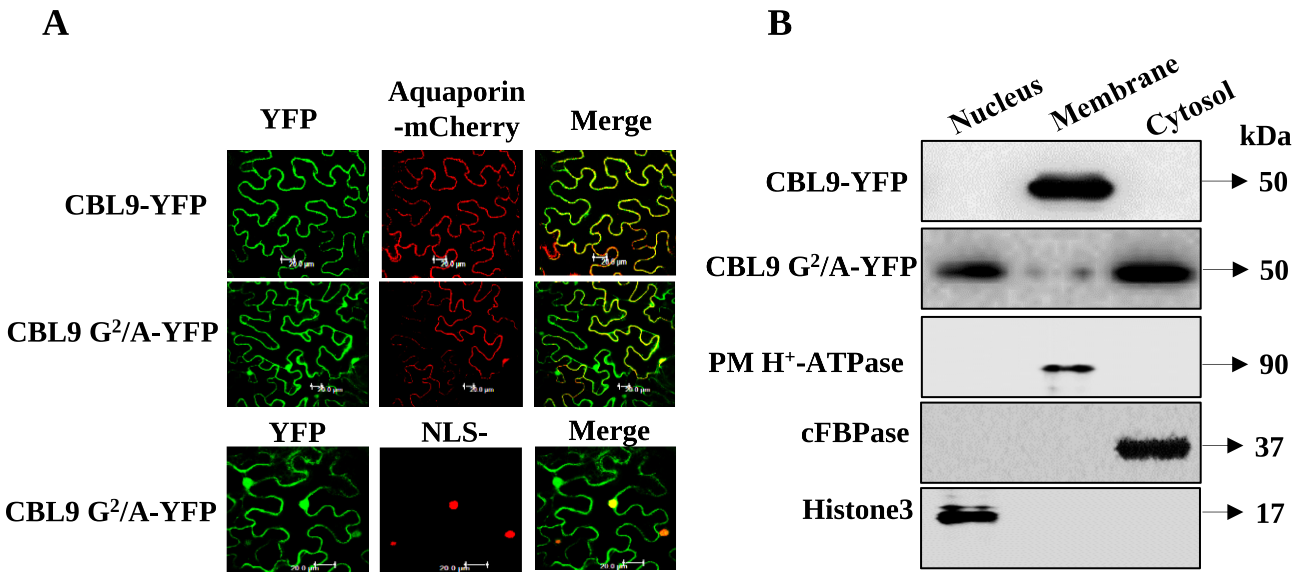

**Figure S6. Localization and subcellular fractionation of CBL9-YFP and CBL9 $G^2$/A-YFP.**

**(A)** CBL9 fused with YFP (green) localizes at Plasma Membrane (PM). Aquaporin-mCherry (red) was used as PM marker and NLS -RFP (red) as nucleus marker. Replacement of second amino acid glycine with alanine ($G^2$/A) in CBL9 alters the localization of CBL9 mostly from PM to cytosol and nucleus with a marginal localization at PM.

**(B)** Immunoblot of subcellular fractions of CBL9-YFP and CBL9 $G^2$/A-YFP exhibited CBL9 is exclusively present in PM while CBL9 $G^2$/A is predominantly present in nucleus and cytosol and marginally in PM fraction. Immunoblot was probed with anti-GFP antibody for CBL9 and CBL9 $G^2$/A. Antibodies against $H^+$-ATPase (plasma membrane), fructose-1,6-bisphosphatase (cFBPase, Cytosolic), and H3 histone (nuclear) were used to detect the markers.

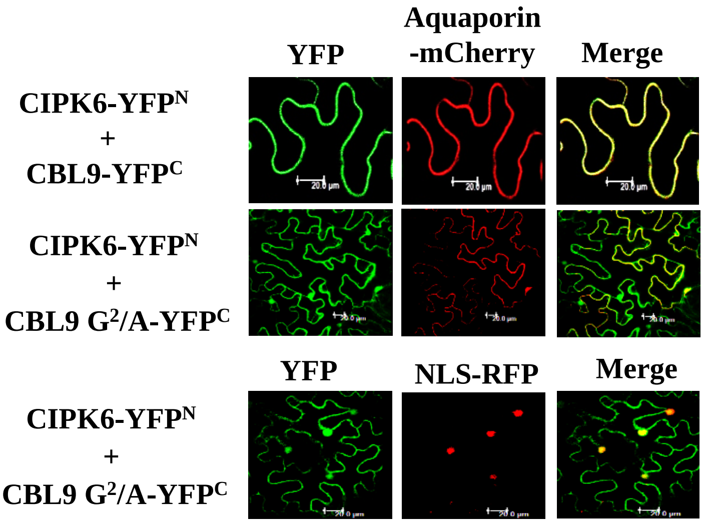

**Figure S7. Interaction of CBL9 G²/A with CIPK6 by BiFC assay.** BiFC showing mutation at myristoylation site (G²/A) of CBL9 alters the localization of CBL9-CIPK6 complex. Aquaporin-mCherry and NLS-RFP were used as Plasma Membrane (PM) and nuclear markers.

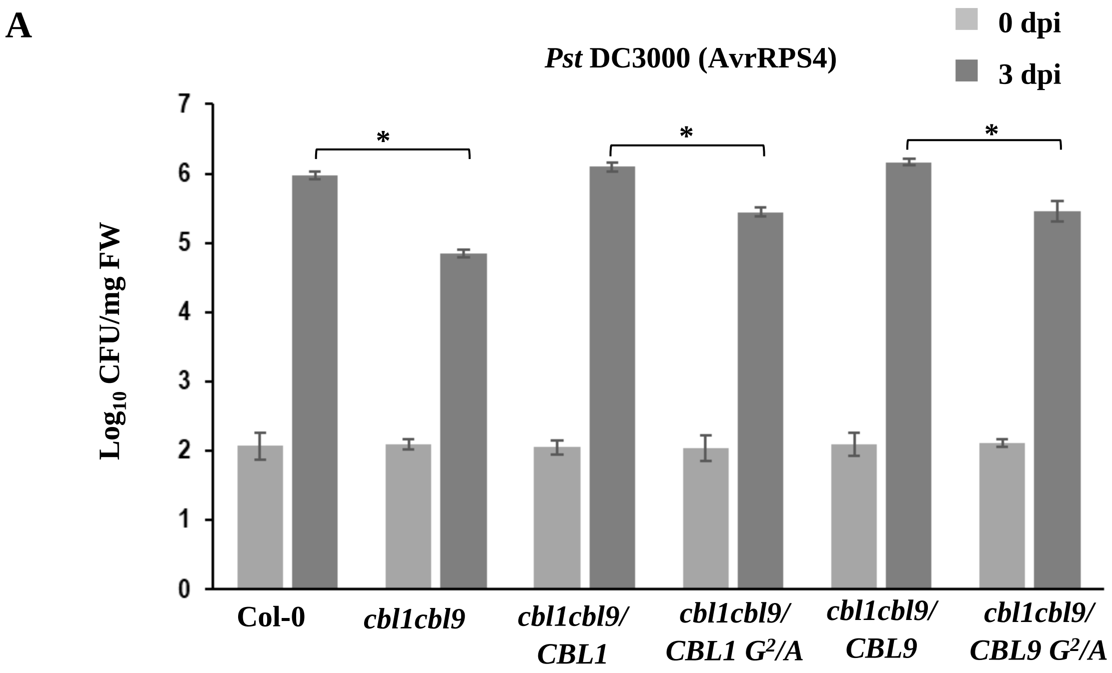

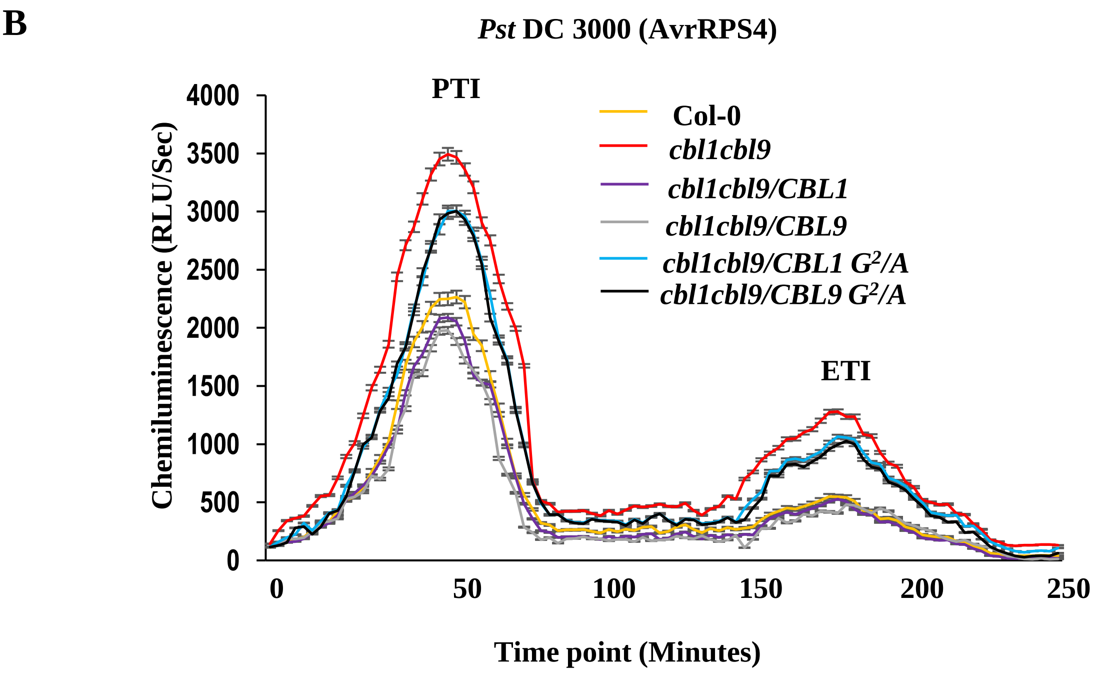

**Figure S8. Altered localization of CBL1/9-CIPK6 complex affects its role in immune response**

**(A)** Bacterial count (cfu/mg FW) was evaluated in in Col-0, *cbl1cbl9*, *cbl1cbl9/CBL1*, *cbl1cbl9/CBL9*, *cbl1cbl9/CBL1 G²/A* and *cbl1cbl9/CBL9 G²/A* plants after infection with *Pst* DC3000 (AvrRPS4) ($OD_{600}$=0.001) at 0 dpi and 3 dpi. The asterisks indicate a significant difference following two-way ANOVA ($\alpha$=0.05).

**(B)** Time-course of ROS production by *in vivo* luminol based assay in response to *Pst* DC3000 (AvrRPS4) ($OD_{600}$ = 0.02) infiltration in Col-0, *cbl1cbl9*, *cbl1cbl9/CBL1*, *cbl1cbl9/CBL9*, *cbl1cbl9/CBL1 G²/A* and *cbl1cbl9/CBL9 G²/A* leaf discs. Results show means (±SE), n>35 leaf discs. ROS peaks of PAMP-triggered and Effector-triggered immune (PTI, ETI) response were shown.

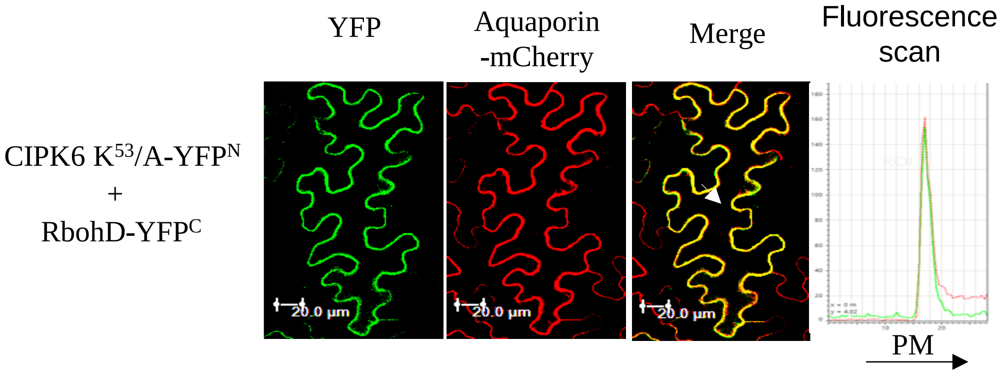

**Figure S9. Interaction of CIPK6 $K^{53}$/A with RbohD by BiFC assay.** CIPK6 $K^{53}$/A-$YFP^{N}$ interacts with RbohD-$YFP^{C}$ at PM *in planta*. Co-localization with PM marker Aquaporin-mCherry was shown.

**A**

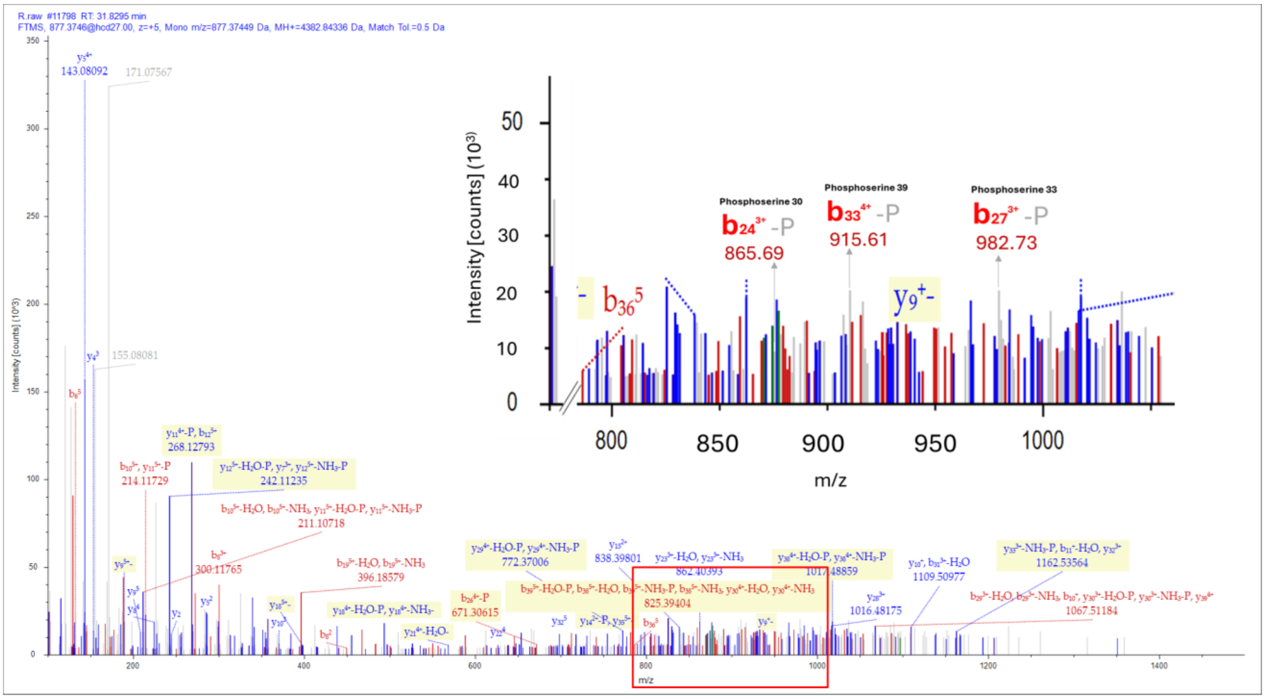

| | Fragment Matches | | | | |
|---|---|---|---|---|---|
| | Value Type: Theo. Mass [Da] | | | | |
| | Ion Series | Phosphorylation Losses | Neutral Losses | Multiple Neutral Losses | Precursor Ions |

| | #1 | b+ | b2+ | b3+ | b4+ | b5+ | Seq. | y+ | y2+ | y3+ | y4+ | y5+ | #2 |
|---|---|---|---|---|---|---|---|---|---|---|---|---|---|
| | 2 | 202.08223 | 101.54475 | 68.03226 | 51.27602 | 41.22227 | S | 4268.82826 | 2134.91777 | 1423.61427 | 1067.96252 | 854.57147 | 39 |
| | 3 | 289.11426 | 145.06077 | 97.04294 | 73.03402 | 58.62867 | S | 4181.79623 | 2091.40175 | 1394.60359 | 1046.20451 | 837.16507 | 38 |
| | 4 | 403.15719 | 202.08223 | 135.05725 | 101.54475 | 81.43726 | N | 4094.76420 | 2047.88574 | 1365.59292 | 1024.44651 | 819.75866 | 37 |
| | 5 | 518.18413 | 259.59570 | 173.39956 | 130.30149 | 104.44265 | D | 3980.72127 | 1990.86428 | 1327.57861 | 995.93578 | 796.95008 | 36 |
| | 6 | 655.24304 | 328.12516 | 219.08587 | 164.56622 | 131.85443 | H | 3865.69433 | 1933.35080 | 1289.23629 | 967.17904 | 773.94469 | 35 |
| | 7 | 784.28564 | 392.64646 | 262.10006 | 196.82687 | 157.66295 | E | 3728.63542 | 1864.82135 | 1243.54999 | 932.91431 | 746.53291 | 34 |
| | 8 | 897.36970 | 449.18849 | 299.79475 | 225.09788 | 180.27976 | L | 3599.59283 | 1800.30005 | 1200.53579 | 900.65366 | 720.72439 | 33 |
| | 9 | 954.39116 | 477.69922 | 318.80191 | 239.35325 | 191.68405 | G | 3486.50876 | 1743.75802 | 1162.84111 | 872.38265 | 698.10757 | 32 |
| | 10 | 1067.47523 | 534.24125 | 356.49659 | 267.62426 | 214.30087 | I | 3429.48730 | 1715.24729 | 1143.83395 | 858.12728 | 686.70328 | 31 |
| | 11 | 1180.55929 | 590.78328 | 394.19128 | 295.89528 | 236.91768 | L | 3316.40323 | 1658.70526 | 1106.13926 | 829.85627 | 664.08647 | 30 |
| | 12 | 1336.66040 | 668.83384 | 446.22499 | 334.92056 | 268.13790 | R | 3203.31917 | 1602.16322 | 1068.44457 | 801.58525 | 641.46966 | 29 |
| | 13 | 1393.68187 | 697.34457 | 465.23214 | 349.17592 | 279.54219 | G | 3047.21806 | 1524.11267 | 1016.41087 | 762.55997 | 610.24943 | 28 |
| | 14 | 1464.71898 | 732.86313 | 488.91118 | 366.93520 | 293.74962 | A | 2990.19660 | 1495.60194 | 997.40372 | 748.30461 | 598.84514 | 27 |
| | 15 | 1578.76191 | 789.88459 | 526.92549 | 395.44593 | 316.55820 | N | 2919.15948 | 1460.08338 | 973.72468 | 730.54533 | 584.63772 | 26 |
| | 16 | 1665.79394 | 833.40061 | 555.93616 | 417.20394 | 333.96461 | S | 2805.11655 | 1403.06192 | 935.71037 | 702.03460 | 561.82913 | 25 |
| | 17 | 1780.82088 | 890.91408 | 594.27848 | 445.96068 | 356.97000 | D | 2718.08453 | 1359.54590 | 906.69969 | 680.27659 | 544.42273 | 24 |
| | 18 | 1881.86856 | 941.43792 | 627.96104 | 471.22260 | 377.17953 | T | 2603.05758 | 1302.03243 | 868.35738 | 651.51985 | 521.41734 | 23 |
| | 19 | 1995.91148 | 998.45938 | 665.97535 | 499.73333 | 399.98812 | N | 2502.00991 | 1251.50859 | 834.67482 | 626.25793 | 501.20780 | 22 |
| | 20 | 2082.94351 | 1041.97539 | 694.98602 | 521.49134 | 417.39452 | S | 2387.96698 | 1194.48713 | 796.66051 | 597.74720 | 478.39922 | 21 |
| | 21 | 2197.97046 | 1099.48887 | 733.32834 | 550.24807 | 440.39991 | D | 2300.93495 | 1150.97111 | 767.64983 | 575.98919 | 460.99281 | 20 |
| | 22 | 2299.01813 | 1150.01271 | 767.01090 | 575.50999 | 460.60945 | T | 2185.90801 | 1093.45764 | 729.30752 | 547.23246 | 437.98742 | 19 |
| **Phosphoserine 30** | 23 | 2428.06073 | 1214.53400 | 810.02509 | 607.77064 | 486.41797 | E | 2084.86033 | 1042.93380 | 695.62496 | 521.97054 | 417.77789 | 18 |
| | 24 | 2595.05909 | 1298.03318 | 865.69121 | 649.52023 | 519.81764 | S-Phosp... | 1955.81774 | 978.41251 | 652.61076 | 489.70989 | 391.96937 | 17 |
| | 25 | 2708.14315 | 1354.57521 | 903.38590 | 677.79125 | 542.43445 | I | 1788.81938 | 894.91333 | 596.94464 | 447.96030 | 358.56970 | 16 |
| **Phosphoserine 33** | 26 | 2779.18026 | 1390.09377 | 927.06494 | 695.55052 | 556.64187 | A | 1675.73531 | 838.37129 | 559.24996 | 419.68929 | 335.95288 | 15 |
| | 27 | 2946.17862 | 1473.59295 | 982.73106 | 737.30011 | 590.04155 | S-Phosp... | 1604.69820 | 802.85274 | 535.57092 | 401.93001 | 321.74546 | 14 |
| | 28 | 3061.20557 | 1531.10642 | 1021.07337 | 766.05685 | 613.04693 | D | 1437.69984 | 719.35356 | 479.90480 | 360.18042 | 288.34579 | 13 |
| | 29 | 3217.30668 | 1609.15698 | 1073.10708 | 805.08213 | 644.26716 | R | 1322.67290 | 661.84009 | 441.56248 | 331.42368 | 265.34040 | 12 |
| | 30 | 3274.32814 | 1637.66771 | 1092.11423 | 819.33749 | 655.67145 | G | 1166.57178 | 583.78953 | 389.52878 | 292.39840 | 234.12018 | 11 |
| | 31 | 3345.36525 | 1673.18627 | 1115.79327 | 837.09677 | 669.87887 | A | 1109.55032 | 555.27880 | 370.52162 | 278.14304 | 222.71589 | 10 |
| **Phosphoserine 39** | 32 | 3492.43367 | 1746.72047 | 1164.81607 | 873.86387 | 699.29256 | F | 1038.51321 | 519.76024 | 346.84259 | 260.38376 | 208.50846 | 9 |
| | 33 | 3659.43203 | 1830.21965 | 1220.48219 | 915.61346 | 732.69223 | S-Phosp... | 891.44479 | 446.22604 | 297.81978 | 223.61666 | 179.09478 | 8 |
| | 34 | 3716.45349 | 1858.73038 | 1239.48935 | 929.86883 | 744.09652 | G | 724.44643 | 362.72686 | 242.15366 | 181.86707 | 145.69511 | 7 |
| | 35 | 3813.50626 | 1907.25677 | 1271.84027 | 954.13202 | 763.50707 | P | 667.42497 | 334.21612 | 223.14651 | 167.61170 | 134.29082 | 6 |
| | 36 | 3926.59032 | 1963.79880 | 1309.53496 | 982.40304 | 786.12389 | L | 570.37221 | 285.68974 | 190.79559 | 143.34851 | 114.88026 | 5 |
| | 37 | 3983.61178 | 1992.30953 | 1328.54211 | 996.65840 | 797.52818 | G | 457.28814 | 229.14771 | 153.10090 | 115.07749 | 92.26345 | 4 |
| | 38 | 4139.71289 | 2070.36009 | 1380.57582 | 1035.68368 | 828.74840 | R | 400.26668 | 200.63698 | 134.09374 | 100.82213 | 80.85916 | 3 |
| | 39 | 4236.76566 | 2118.88647 | 1412.92674 | 1059.94687 | 848.15895 | P | 244.16557 | 122.58642 | 82.06004 | 61.79685 | 49.63893 | 2 |
| | 40 | | | | | | K | 147.11280 | 74.06004 | 49.70912 | 37.53366 | 30.22838 | 1 |

**B**

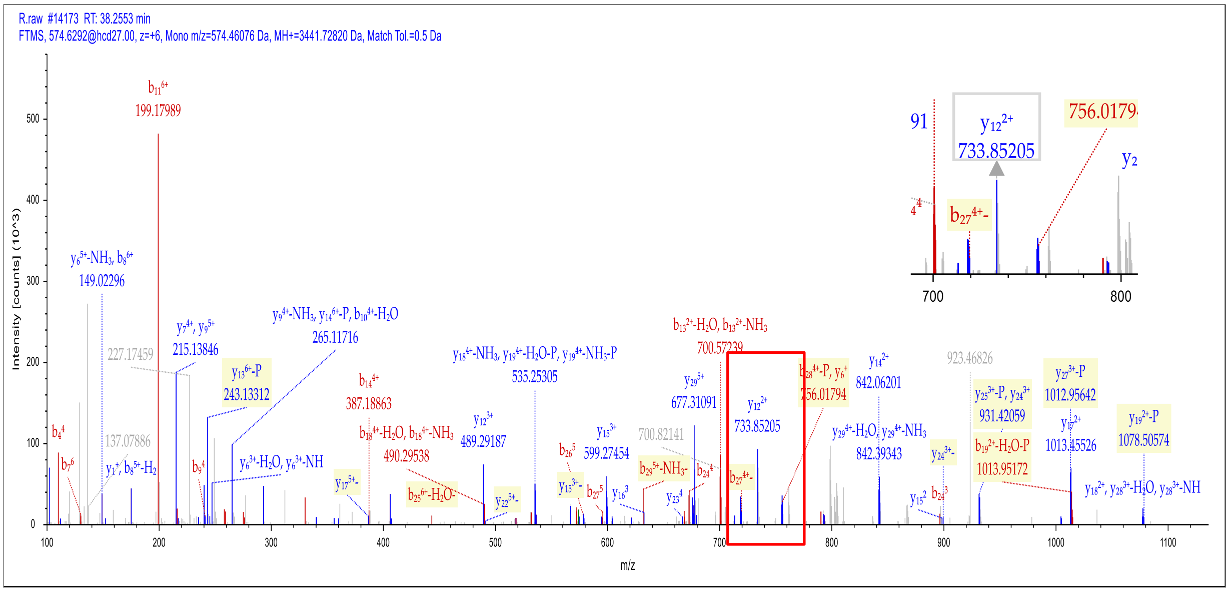

Fragment Matches

Value Type: Theo. Mass [Da]

Ion Series | Phosphorylation Losses | Neutral Losses | Multiple Neutral Losses | Precursor Ions

| #1 | b+ | b2+ | b3+ | b4+ | b5+ | b6+ | Seq. | y+ | y2+ | y3+ | y4+ | y5+ | y6+ | #2 |
|---|---|---|---|---|---|---|---|---|---|---|---|---|---|---|
| 1 | 58.02874 | 29.51801 | 20.01443 | 15.26264 | 12.41157 | 10.51085 | G | | | | | | | 30 |
| 2 | 195.08765 | 98.04746 | 65.70074 | 49.52737 | 39.82335 | 33.35401 | H | 3384.70868 | 1692.85798 | 1128.90774 | 846.93263 | 677.74756 | 564.95751 | 29 |
| 3 | 308.17172 | 154.58950 | 103.39542 | 77.79839 | 62.44016 | 52.20135 | L | 3247.64977 | 1624.32852 | 1083.22144 | 812.66790 | 650.33577 | 542.11436 | 28 |
| 4 | 437.21431 | 219.11079 | 146.40962 | 110.05903 | 88.24868 | 73.70845 | E | 3134.56570 | 1567.78649 | 1045.52675 | 784.39688 | 627.71896 | 523.26701 | 27 |
| 5 | 552.24125 | 276.62426 | 184.75194 | 138.81577 | 111.25407 | 92.87961 | D | 3005.52311 | 1503.26519 | 1002.51255 | 752.13623 | 601.91044 | 501.75992 | 26 |
| 6 | 649.29402 | 325.15065 | 217.10286 | 163.07896 | 130.66462 | 109.05507 | P | 2890.49617 | 1445.75172 | 964.17024 | 723.37950 | 578.90505 | 482.58876 | 25 |
| 7 | 778.33661 | 389.67194 | 260.11705 | 195.33961 | 156.47314 | 130.56217 | E | 2793.44340 | 1397.22534 | 931.81932 | 699.11631 | 559.49450 | 466.41330 | 24 |
| 8 | 891.42067 | 446.21397 | 297.81174 | 223.61063 | 179.08996 | 149.40951 | L | 2664.40081 | 1332.70404 | 888.80512 | 666.85566 | 533.68598 | 444.90620 | 23 |
| 9 | 962.45779 | 481.73253 | 321.49078 | 241.36990 | 193.29738 | 161.24903 | A | 2551.31675 | 1276.16201 | 851.11043 | 638.58464 | 511.06917 | 426.05885 | 22 |
| 10 | 1075.54185 | 538.27456 | 359.18547 | 269.64092 | 215.91419 | 180.09637 | L | 2480.27963 | 1240.64345 | 827.43139 | 620.82537 | 496.86175 | 414.21934 | 21 |
| 11 | 1188.62591 | 594.81660 | 396.88016 | 297.91194 | 238.53100 | 198.94372 | L | 2367.19557 | 1184.10142 | 789.73671 | 592.55435 | 474.24493 | 395.37199 | 20 |
| 12 | 1289.67359 | 645.34043 | 430.56272 | 323.17386 | 258.74054 | 215.78500 | T | 2254.11150 | 1127.55939 | 752.04202 | 564.28333 | 451.62812 | 376.52465 | 19 |
| 13 | 1417.76856 | 709.38792 | 473.26104 | 355.19760 | 284.35953 | 237.13416 | K | 2153.06383 | 1077.03555 | 718.35946 | 539.02141 | 431.41859 | 359.68337 | 18 |
| 14 | 1545.86352 | 773.43540 | 515.95936 | 387.22134 | 309.97853 | 258.48332 | K | 2024.96886 | 1012.98807 | 675.66114 | 506.99767 | 405.79959 | 338.33421 | 17 |
| 15 | 1646.91120 | 823.95924 | 549.64192 | 412.48326 | 330.18806 | 275.32460 | T | 1896.87390 | 948.94059 | 632.96282 | 474.97393 | 380.18060 | 316.98505 | 16 |
| 16 | 1759.99526 | 880.50127 | 587.33660 | 440.75427 | 352.80487 | 294.17194 | L | 1795.82622 | 898.41675 | 599.28026 | 449.71201 | 359.97107 | 300.14377 | 15 |
| 17 | 1889.03785 | 945.02257 | 630.35080 | 473.01492 | 378.61339 | 315.67904 | E | 1682.74216 | 841.87472 | 561.58557 | 421.44100 | 337.35425 | 281.29642 | 14 |
| 18 | 1976.06988 | 988.53858 | 659.36148 | 494.77293 | 396.01980 | 330.18438 | S | 1553.69956 | 777.35342 | 518.57137 | 389.18035 | 311.54573 | 259.78932 | 13 |
| 19 | 2143.06824 | 1072.03776 | 715.02760 | 536.52252 | 429.41947 | 358.01744 | S-Phosp... | 1466.66754 | 733.83741 | 489.56070 | 367.42234 | 294.13933 | 245.28399 | 12 |
| 20 | 2256.15231 | 1128.57979 | 752.72229 | 564.79353 | 452.03628 | 376.86478 | L | 1299.66918 | 650.33823 | 433.89458 | 325.67275 | 260.73966 | 217.45093 | 11 |
| 21 | 2370.19523 | 1185.60125 | 790.73660 | 593.30427 | 474.84487 | 395.87194 | N | 1186.58511 | 593.79619 | 396.19989 | 297.40174 | 238.12284 | 198.60358 | 10 |
| 22 | 2484.23816 | 1242.62272 | 828.75090 | 621.81500 | 497.65345 | 414.87909 | N | 1072.54218 | 536.77473 | 358.18558 | 268.89100 | 215.31426 | 179.59643 | 9 |
| 23 | 2585.28584 | 1293.14656 | 862.43346 | 647.07692 | 517.86299 | 431.72037 | T | 958.49926 | 479.75327 | 320.17127 | 240.38027 | 192.50567 | 160.58927 | 8 |
| 24 | 2686.33352 | 1343.67040 | 896.11602 | 672.33884 | 538.07252 | 448.56165 | T | 857.45158 | 429.22943 | 286.48871 | 215.11835 | 172.29614 | 143.74799 | 7 |
| 25 | 2773.36555 | 1387.18641 | 925.12670 | 694.09684 | 555.47893 | 463.06699 | S | 756.40390 | 378.70559 | 252.80615 | 189.85643 | 152.08660 | 126.90671 | 6 |
| 26 | 2886.44961 | 1443.72844 | 962.82139 | 722.36786 | 578.09574 | 481.91433 | L | 669.37187 | 335.18957 | 223.79548 | 168.09843 | 134.68020 | 112.40138 | 5 |
| 27 | 2973.48164 | 1487.24446 | 991.83206 | 744.12587 | 595.50215 | 496.41967 | S | 556.28781 | 278.64754 | 186.10079 | 139.82741 | 112.06338 | 93.55403 | 4 |
| 28 | 3120.55005 | 1560.77866 | 1040.85487 | 780.89297 | 624.91583 | 520.93107 | F | 469.25578 | 235.13153 | 157.09011 | 118.06940 | 94.65698 | 79.04869 | 3 |
| 29 | 3267.61847 | 1634.31287 | 1089.87767 | 817.66007 | 654.32951 | 545.44247 | F | 322.18737 | 161.59732 | 108.06731 | 81.30230 | 65.24329 | 54.53729 | 2 |
| 30 | | | | | | | R | 175.11895 | 88.06311 | 59.04450 | 44.53520 | 35.82961 | 30.02589 | 1 |

**Phosphoserine 119**

**Figure S10. Mass spectrometry analysis of phosphorylated RbohD. (A-B)** Mass spectrometry analysis of CIPK6-phosphorylated RbohD-N. The spectra of phosphorylated serine residues at 30, 33, 39, 119 of RbohD and its b and y value were shown in box .

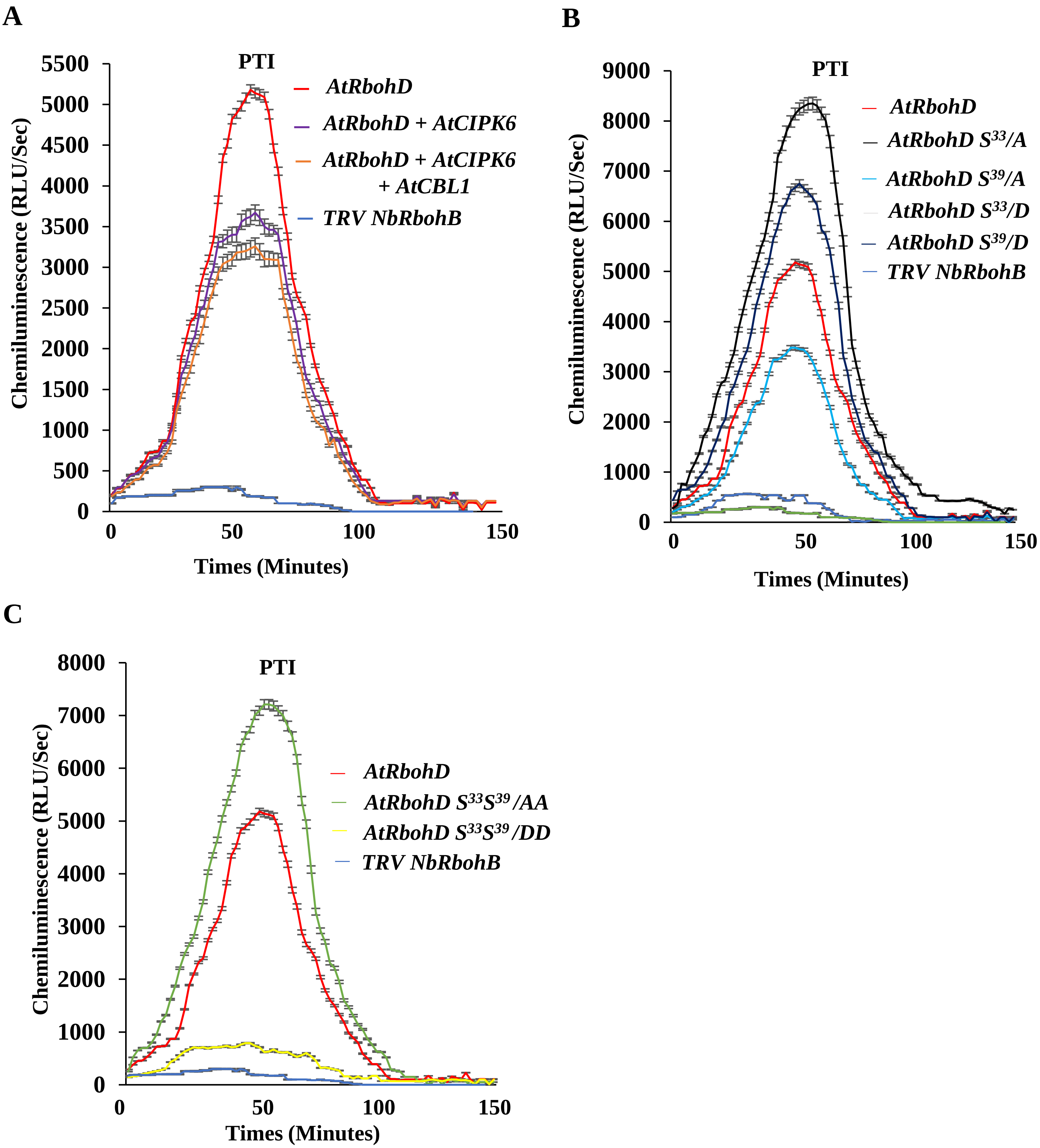

**Figure S11. Time-course of luminol enhanced chemiluminiscence asaay of *in vivo* ROS production in *N. benthamiana TRV-NbRbohB* silenced plants. (A-C)** The proteins mentioned in each figures were transiently expressed in *N. benthamiana TRV-NbRbohB* silenced plants. Relative luminescence unit per second (RLU/sec) was measured in response to *Pst* DC 3000 ($OD_{600}$ = 0.1) inoculation. Results show means (±SE), n>35 leaf discs. Box plots of the maximum values were presented in Fig. 6.

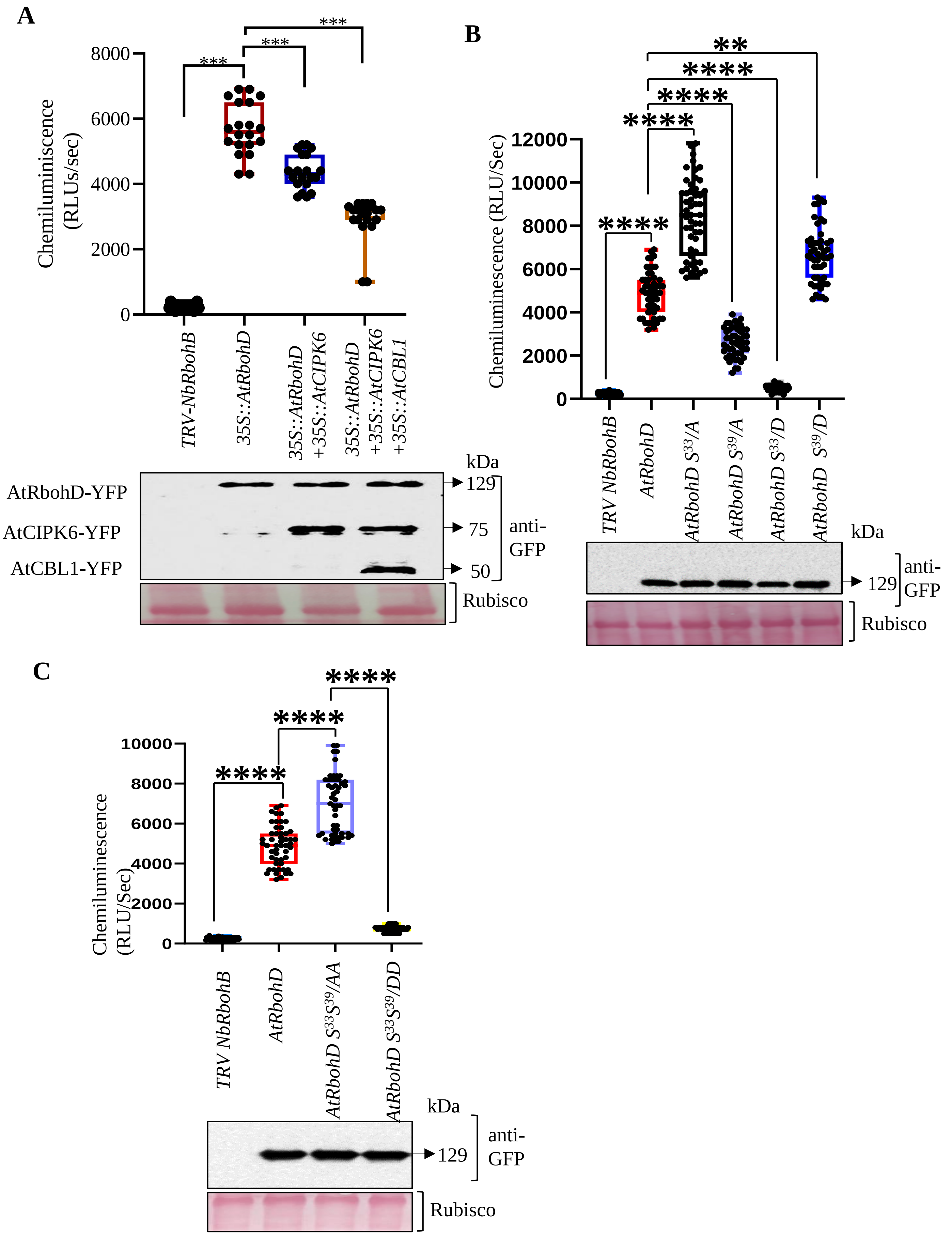

A
Chemiluminiscence (RLUs/sec)
8000
6000
4000
2000
0
***
***
***
TRV-NbRbohB
35S::AtRbohD
35S::AtRbohD +35S::AtCIPK6
35S::AtRbohD +35S::AtCIPK6 +35S::AtCBL1
kDa
AtRbohD-YFP
AtCIPK6-YFP
AtCBL1-YFP
129
75
50
anti-GFP
Rubisco
B
Chemiluminescence (RLU/Sec)
12000
10000
8000
6000
4000
2000
0
**
****
****
****
****
TRV NbRbohB
AtRbohD
AtRbohD S33/A
AtRbohD S39/A
AtRbohD S33/D
AtRbohD S39/D
kDa
129
anti-GFP
Rubisco
C
Chemiluminescence (RLU/Sec)
10000
8000
6000
4000
2000
0
****
****
****
TRV NbRbohB
AtRbohD
AtRbohD S33S39/AA
AtRbohD S33S39/DD
kDa
129
anti-GFP
Rubisco

**Figure S12. CIPK6 and its interacting CBLs negatively regulates the ROS producing activity of RbohD.**

(A) Virus induced gene silencing (VIGS) of *NbRbohB* (*TRV-NbRbohB*) in *Nicotiana* significantly reduced the ROS producing activity of NbRbohB against *Pst* infection. Transient expression of AtRbohD protein in *TRV-NbRbohB* plants restored the ROS production whereas the transient expression of AtRbohD with AtCIPK6, and AtCBL1 moderately reduced the ROS production against *Pst* infection. Box-Plot showing the plot of maximum values of ROS obtained during PTI. $n > 35$ for ROS burst. The asterisks indicate a significant difference analysed through two-way ANOVA ($\alpha$=0.05). Equal expression of AtRbohD, AtCIPK6 and AtCBL1 was detected by anti-GFP immunoblotting in *TRV-NbRbohB* plants.

(B) Single Phosphonull AtRbohD $S^{33}$/A showed more ROS production as compared to AtRbohD $S^{39}$/A and AtRbohD while single phsphomimetic AtRbohD $S^{33}$/D abolished the ROS production as compared to AtRbohD $S^{39}$/D and AtRbohD after *Pst* infection. Box-Plot showing the plot of maximum values of ROS attained during PTI. An immunoblot prepared with anti-GFP Ab demonstrates the transient and equal abundance of AtRbohD, AtRbohD $S^{33}$/A, AtRbohD $S^{39}$/A, AtRbohD $S^{33}$/D and AtRbohD $S^{39}$/D in *NbRbohB* silenced *Nicotiana* leaves.

(C) After *Pst* inoculation double Phosphonull AtRbohD $S^{33}S^{39}$/AA exhibited more ROS production as compared to AtRbohD while AtRbohD $S^{33}S^{39}$/DD showed diminished ROS production. Box-Plot showing the plot of maximum values of ROS attained during PTI. Immunoblot developed through anti-GFP Ab showing the transient and equal expression of AtRbohD, AtRbohD $S^{33}S^{39}$/AA and AtRbohD $S^{33}S^{39}$/DD in *NbRbohB* silenced Nicotiana leaves.

A

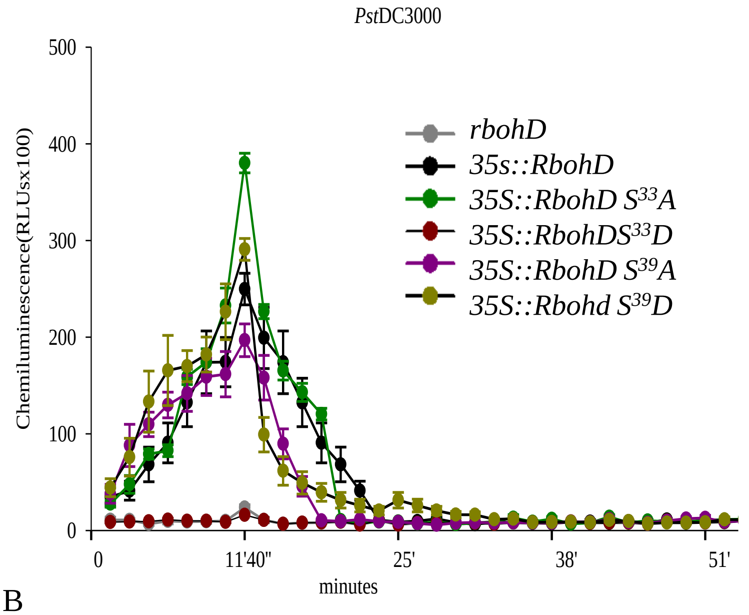

B

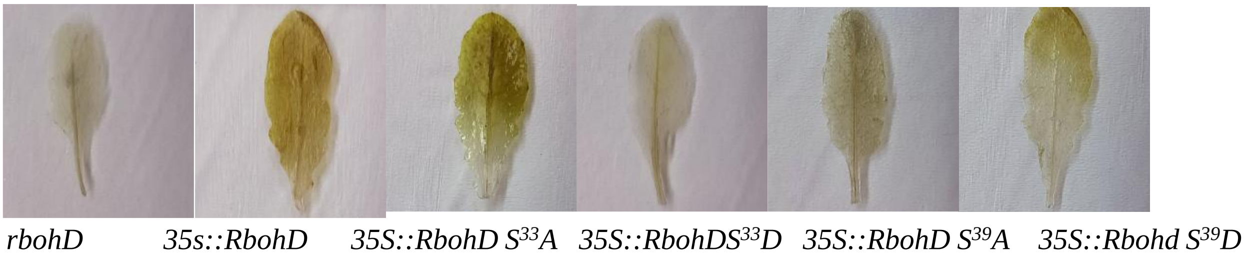

C

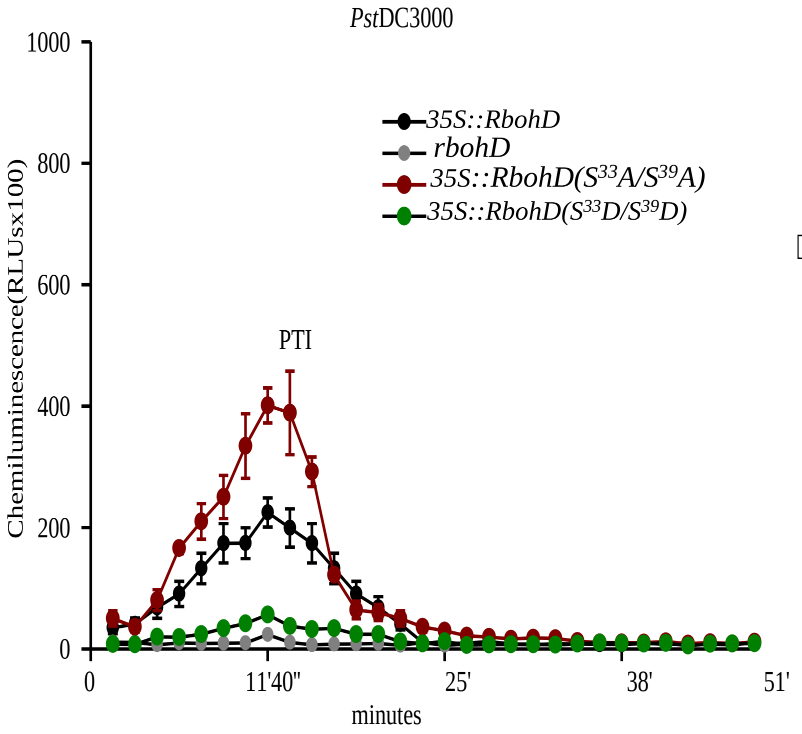

D

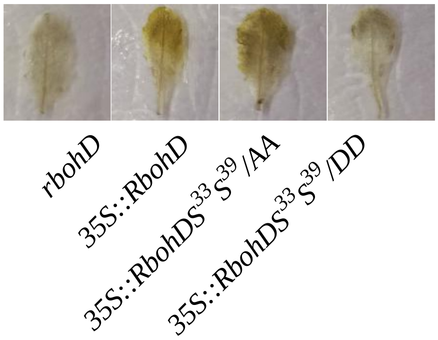

DAB stained leaves after pstDC3000 infiltration

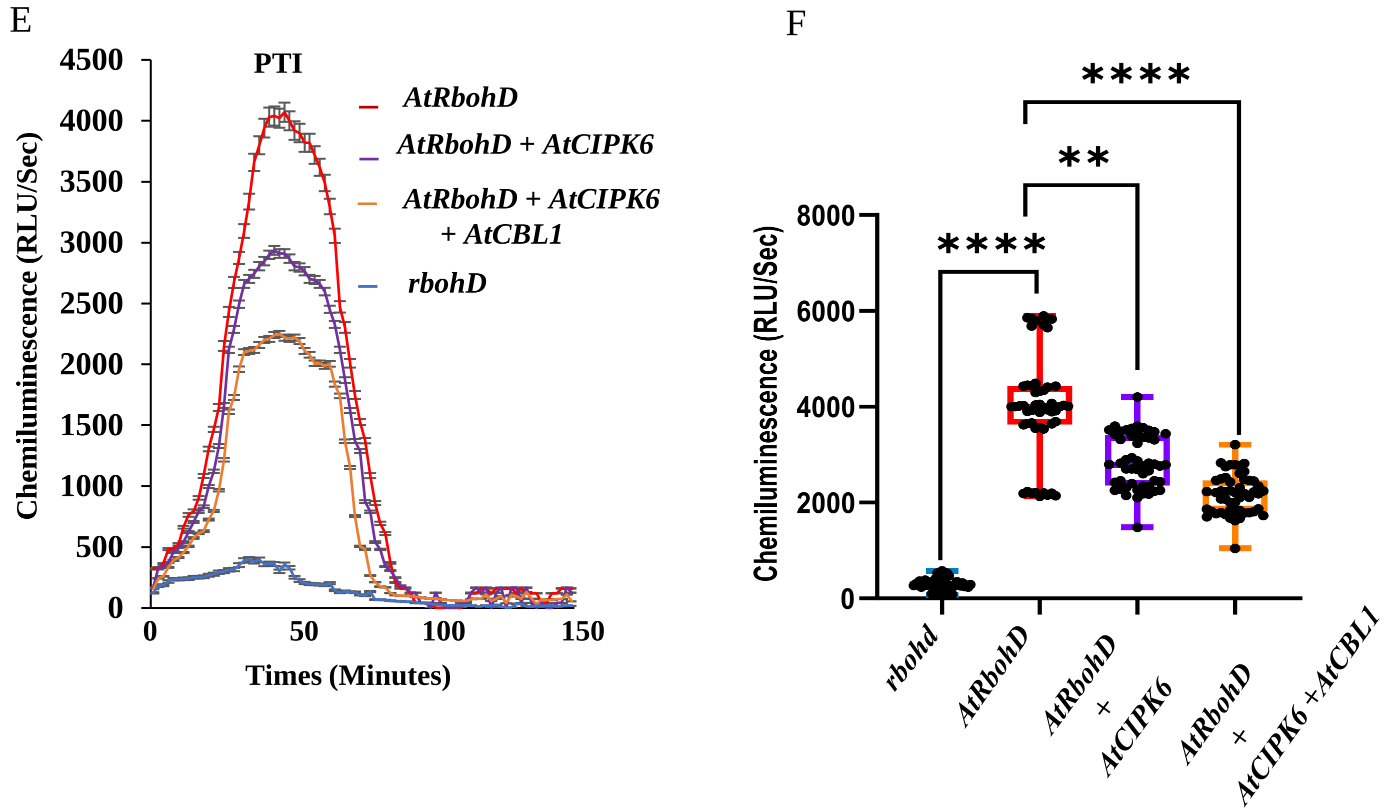

**Figure S13.** ***In vivo*** **luminol based ROS measurement in Arabidopsis.**

(A) Time course of ROS production in *Arabidopsis* plants, over expressing *RbohD*, *RbohD*($S^{33}A$), *RbohD*($S^{39}A$), *RbohD*($S^{33}S^{39}/AA$), *RbohD*($S^{33}D$), *RbohD*($S^{39}D$), *RbohD*($S^{33}S^{39}/DD$), and *rbohD* mutants. (B) DAB stained Arabidopsis leaves. (C) ROS production in *Arabidopsis* plants, over expressing *RbohD*, *RbohD*($S^{33}S^{39}/AA$), *RbohD*($S^{33}S^{39}/DD$), and *rbohD* mutants. Relative luminescence unit per second (RLU/sec) was measured in response to *Pst* DC 3000 ($OD_{600}$ = 0.1) inoculation. Results show means (±SE), n=24 leaf discs. (B) DAB stained Arabidopsis leaves. (D) DAB stained leaves for the indicated genotypes. (E) Time course of ROS generation was done in *rbohD* mutants, *35S::RbohD-GFP*, CIPK6 overexpressing plants in the background of 35S::RbohD and CIPK6 and CBL1 overexpressing in the 35S::RbohD background. (F) Box-Plot showing the plot of maximum values of ROS attained during PTI. $n > 35$ for ROS burst. The asterisks indicate significance as obtained through two-way ANOVA (α=0.05).

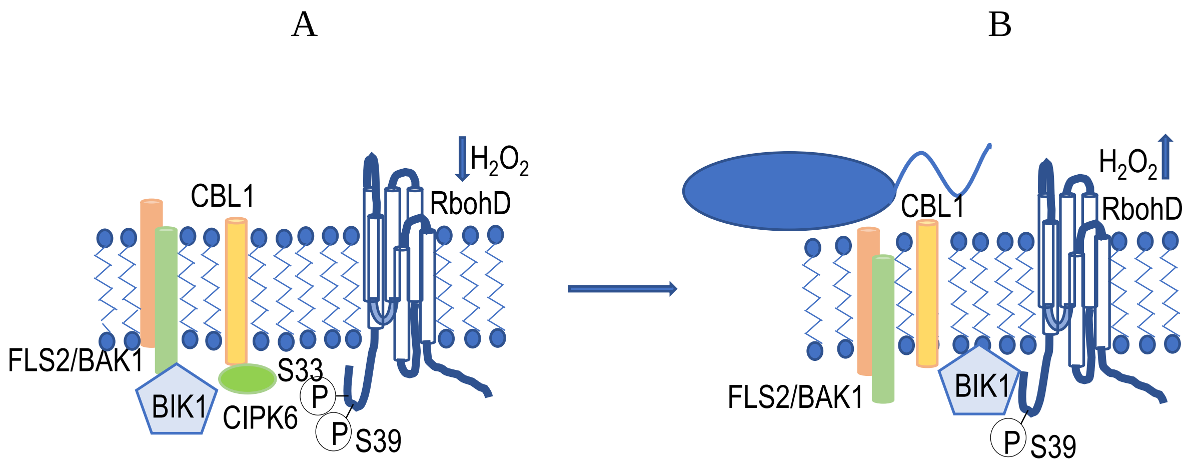

**Figure S14:** Proposed model illustrating the positive and negative regulation of RbohD-mediated ROS generation through the BIK1 and the CBL1/9-CIPK6 module in Arabidopsis. **(A)** In the absence of *Pst*DC3000, the CIPK6 level would be high, allowing it to interact with CBL1/9 and phosphorylate RbohD at $S^{33}$ and $S^{39}$ residues. Since $S^{33}$ is the primary serine site phosphorylated by CIPK6, this modification leads to the negative regulation of ROS generation through RbohD. **(B)** In the presence of *Pst*DC3000, CIPK6 levels decrease rapidly. Simultaneously, the interaction of FLS2 with BAK1 initiates signaling, leading to BAK1 autophosphorylation and subsequent transphosphorylation of the S39 residue of RbohD through BIK1. This modification enhances ROS generation, while the reduced expression of CIPK6 prevents its phosphorylation activity.

**Supplementary Table 1 : Primers list used in this study**

| S.N. | Name of primer | Sequence of primers |
|---|---|---|
| 1 | GKBP_RP | 5’-ATCAGATTGTCGTTTCCCGC-3’ |
| 2 | LBb1.3 | 5'-ATTTTGCCGATTTCGGAAC-3' |
| 3 | CIPK6 GTY_F | 5'- CACCATGGTCGGAGCAAAACCGGTGGAG -3' |
| 4 | CIPK6 GTY_R SCR | 5'- AGCAGGTGTAGAGGTCCAG -3' |
| 5 | CIPK6_F (BamH1) | 5'- CGGGATCCATGGTCGGAGCAAAACCG-3' |
| 6 | CIKP6_R (Xho1) SCR | 5'-GCGTATCTCGAGAGCAGGTGTAGAGGTCCAG-3' |
| 7 | CIPK6-N-ter GTY_R | 5'-CACTGGTTCGTTTCTTGATCTCGT-3' |
| 8 | CIPK6-N-ter_R (EcoR1) SCR | 5'-GAATTCCACTGGTTCGTTTCTTGATCTCGT -3' |
| 9 | CIPK6 $K^{53}$/A SDM_F | 5'- AAGCGTGGCGATGGCCGTCGTCGGAAAAGG -3' |
| 10 | CIPK6 $K^{53}$/A SDM_R | 5'- CTCTTTTCCGACGACGGCCATCGCCACGCTT-3' |
| 11 | CIPK6 $T^{182}$/D SDM _F | 5'GACGGACTTCTTCATGATACTTGTGGAACTCC -3' |
| 12 | CIPK6 $T^{182}$/D SDM _R | 5'- GGAGTTCCACAAGTATCATGAAGAAGTCCGTC -3' |
| 13 | CBL1 GTY_F | 5'-CACCATGGGCTGCTTCCACTCAAAG-3' |
| 14 | CBL1 GTY_R SCR | 5'-TGTGGCAATCTCATCGA-3' |
| 15 | CBL9 GTY_F | 5'-CACCATGGGTTGTTTCCATTCCACG-3' |
| 16 | CBL9 GTY_R SCR | 5'-CGTCGCAATCTCGTCCA-3' |
| 17 | CBL1_F (BamH1) | 5'- CGGGATCCATGGGCTGCTTCCACTCAAAG -3' |
| 18 | CBL1_R (Xho1) SCR | 5'- CCGCTCGACTGTGGCAATCTCATCG-3' |
| 19 | CBL9_F (BamH1) | 5'- CGGGATCCATGGGTTGTTTCCATTCCACG -3' |
| 20 | CBL9_R (Sal1)SCR | 5'- CCGGTCGACCGTCGCAATCTCGTCC -3' |
| 21 | CBL1 $G^{2}$/A SDM_F | 5'-CCCTTCACCATGGCTTGCTTCCACTCAAA-3' |
| 22 | CBL1 $G^{2}$/A SDM_R | 5'-TTTGAGTGGAAGCAAGCCATGGTGAAGGG -3' |
| 23 | CBL9 $G^{2}$/A SDM_F | 5'-GCCCCCTTCACCATGGCTTGTTTCCATTCCACG -3' |
| 24 | CBL9 $G^{2}$/A SDM_R | 5'- CGTGGAATGGAAACAAGCCATGGTGAAGGGGGC -3' |
| 25 | RbohD GTY_F | 5'-CACCATGAAAATGAGACGAGGC-3' |
| 26 | RbohD GTY_R | 5'- GAAGTTCTCTTTGTGGAAGTCAAAC -3' |
| 27 | RbohD N-ter GTY_R SCR | 5'-TCTCTGCCAATTGTCAAGTATG-3' |
| 28 | RbohD _F (Nde1) | 5'- GGACTCCATATGAAAATGAGACGAGGC -3' |
| 29 | RbohD -N __R (ECoR1) SCR | 5'-ATACGGAATTCTCTCTGCCAATTGTCAAGTATG -3' |
| 30 | M13_F | 5'- CCCAGTCACGACGTTGTAAAACG -3’ |

| | | |
|---|---|---|
| 31 | M13_R (pGEMT) | 5'- AGCGGATAACAATTTCACACAGG -3' |
| 32 | M13_R (pENTER) | 5'-CAGGAAACAGCTATGAC -3' |
| 33 | pGEX4T2 sequencing _F primer | 5’- GTATTGAAGCTATCCCACAAATTG -3’ |
| 34 | pGEX4T2 sequencing _R primer | 5'- TTGTCTGCTCCCGGCATCCGCTTA -3' |
| 35 | 35s Promoter Uni_F | 5'- CACTATCCTTCGCAAGACCCTTCC -3' |
| 36 | YFP Uni_R | 5'- CTTGAAGAAGATGGTGCGCTCCTGGAC -3' |
| 37 | YFP_F1 | 5'- ATGAAGCAGCACGACTTCTTCAAGT -3' |
| 38 | YFP_F2 | 5'- ATGGTGAGCAAGGGCGAGGAGCTG -3' |
| 39 | YFPc_R | 5'- CTGGTAGTGGTCGGCGAGCTGCACGCT-3' |
| 40 | RbohD $S^{30}$/D_F | 5'- GGACACGGAGGACATCGCTAGCGACCG-3' |
| 41 | RbohD $S^{30}$/D_R | 5'- CGGTCGCTAGCGATGTCCTCCGTGTCC-3' |
| 42 | RbohD $S^{33}$/D_F | 5'- GGACACGGAGAGCATCGCTGACGACCG-3' |
| 43 | RbohD $S^{33}$/D_R | 5'- CGGTCGTCAGCGATGCTCTCCGTGTCC-3' |
| 44 | RbohD $S^{39}$/D_F | 5'- CGTGGTGCCTTTGACGGTCCGCTTG-3' |
| 45 | RbohD $S^{39}$/D_R | 5'- CAAGCGGACCGTCAAAGGCACCACG-3' |
| 46 | RbohD $S^{119}$/D_F | 5'- GAAGACTCTCGAGAGCGACCTCAACAACACCAC-3' |
| 47 | RbohD $S^{119}$/D_R | 5'- GTGGTGTTGTTGAGGTCGCTCTCGAGAGTCTTC-3' |
| 48 | RbohD $S^{33}S^{39}$/DD_F | 5'- ACGGAGAGCATCGCTGACGACCGTGGTGCCTTTGACGGTCCG-3' |
| 49 | RbohD $S^{33}S^{39}$/DD_R | 5'- CGGACCGTCAAAGGCACCACGGTCGTCAGCGATGCTCTCCGT-3' |
| 50 | RbohD $S^{30}$/A_F | 5'- GGACACGGAGGCCATCGCTAGCGACCG-3' |
| 51 | RbohD $S^{30}$/A_R | 5'- CGGTCGCTAGCGATGGCCTCCGTGTCC-3' |
| 52 | RbohD $S^{33}$/A_F | 5'- GGACACGGAGAGCATCGCTGCCGACCG-3' |
| 53 | RbohD $S^{33}$/_R | 5'- CGGTCGGCAGCGATGCTCTCCGTGTCC-3' |
| 54 | RbohD $S^{39}$/A_F | 5'- CGTGGTGCCTTTGCCGGTCCGCTTG-3' |
| 55 | RbohD $S^{39}$/A_R | 5'- CAAGCGGACCGGCAAAGGCACCACG-3' |
| 56 | RbohD $S^{119}$/A_F | 5'- GAAGACTCTCGAGAGCGCCCTCAACAACACCAC-3' |
| 57 | RbohD $S^{119}$/A_R | 5'- GTGGTGTTGTTGAGGGCGCTCTCGAGAGTCTTC-3' |
| 58 | RbohD $S^{33}S^{39}$/AA_F | 5'- ACGGAGAGCATCGCTGCCGACCGTGGTGCCTTTGCCGGTCCG-3' |
| 59 | RbohD $S^{33}S^{39}$/AA_R | 5'- CGGACCGGCAAAGGCACCACGGTCGGCAGCGATGCTCTCCGT-3' |
| 60 | NtRbohB_F (BamH1) | 5'- GGATCCAATCATCATCCGCACCACCATCAC-3' |
| 61 | NtRbohB_R (ECoR1) | 5'- GAATTCCGTAGGCATCATCATTGGAC-3' |
